# Domain consolidation in Bacterial 50S assembly revealed by Anti-Sense Oligonucleotide Probing

**DOI:** 10.1101/2024.05.08.593220

**Authors:** Kai Sheng, Xiyu Dong, Sriram Aiyer, Joan Lee, Selena Djordjevic Marquardt, Dmitry Lyumkis, James R. Williamson

## Abstract

Investigating the intricate and rapid folding kinetics of large RNA-protein complexes (RNPs), like the bacterial ribosome, remains a formidable challenge in structural biology. Previous genetic approaches to probe assembly have focused on modulating the expression of either r-proteins or assembly factors. Here, anti-sense oligonucleotides (ASOs) were used to disrupt native RNA/RNA and RNA/protein interactions, in order to generate novel folding intermediates. In an *in vitro* co-transcriptional assembly assay, 8 assembly inhibitor ASOs were identified. Using cryo-electron microscopy, 38 new intermediate structures were determined resulting from the specific inhibitions. In particular a novel intermediate class provided compelling evidence of independent rRNA domain folding before proper interdomain docking. Three PNAs targeting domain-I of 23S- rRNA further subdivided the previously identified assembly core into smaller blocks representing the earliest steps in assembly. The resulting assembly graph reveals template-directed RNA foldon docking and domain consolidation, which provides a hierarchical view of the RNP assembly process.

## Introduction

Ribosome biogenesis is efficient in cell, and in *E. coli*, the assembly time of a 70S ribosome is on the order of ∼2 minutes.^1,2^ In recent years, the mechanistic understanding of ribosome assembly has advanced to include a structural framework for sequential and cooperative domain assembly. While heterogeneous reconstruction methods that can deconvolute ensembles of assembly intermediates are developing, studies thus far have primarily focused on genetic manipulation of ribosomal proteins (r-proteins)^3^ or assembly factors^4–9^ in order to induce accumulation of intermediates due to an assembly defect. Nikolay *et al.* also used genetically tagged assembly factor to enrich ObgE-bound intermediates^10^. Similarly, Zhang *et al.* incubated EngA with chemically disrupted 50S to capture an EngA bound intermediate^11^. In addition, *in vitro* reconstitution has provided a key connection between the classic assembly maps and the intermediates observed *in vitro* and in cells^6,12^.

About half of the mass of ribosome is ribosomal RNA (rRNA). The folding of 16S rRNA has been explored by hydroxyl radical probing both *in vitro*^13^ and *in vivo*^14^ led by Woodson’s lab, offering valuable RNA folding kinetics for 30S biogenesis. However, the details of folding of 50S rRNA (23S and 5S) have yet to be elucidated. There are more than 130 RNA helices in 23S and 5S defined by phylogenetically conserved secondary structure.^15^ Among these helices in the bacterial large ribosomal subunit (LSU), there is an intricate tertiary interaction network including hundreds of A-minor motif interactions, long-distance base pairing, magnesium bridges, to name a few types. Recent study from Woodson’s lab using time-resolved DMS-MAP showed late stage rRNA consolidation events in 50S.^16^ However, few early events were observed due to the fast kinetics and small population of early assembly intermediates. The order of formation of most of the interactions that drive the early assembly process remains largely unknown. One concrete example is the smallest assembly core composed of ∼600 nucleotides of 23S rRNA, reported by three independent studies^6,7,12^ , which has not yet been fully dissected to reveal the earliest steps in 23S rRNA folding.

While previous perturbations of r-protein expression or deletion of assembly factors have proven to be extremely valuable, it is generally difficult to anticipate the consequence of a particular perturbation. Thanks to the sequence specificity of nucleic acid base pairing, complementary oligonucleotides and their analogs have been used for site-specific probing RNA structure and folding kinetics for decades,^17,18^ . However, these studies only focused on smaller RNAs or simpler RNPs using biochemical assays. Hybridization probing has not been widely applied to ribosome assembly, and the structures of trapped intermediates has not been elucidated.

**I**ntegrated **S**ynthesis of rRNA, ribosome **A**ssembly and **T**ranslation assay^19^, iSAT assay, is a powerful platform for monitoring *in vitro* assembly of bacterial ribosome (**Fig.1 a**), and we have shown that the process generally resembles near-physiological assembly.^6^ The iSAT system offers the opportunity to probe ribosome assembly by introducing site specific oligonucleotide probes that have the opportunity to kinetically compete with assembly and the potential to arrest assembly, thereby revealing the intermediates resulting from this specific perturbation. Inhibition of assembly is observed by the effect on translation of a GFP reporter construct, and the stalled intermediates can be isolated for structural characterization using cryo-EM. Using the iSAT system as a platform, we screened a small targeted oligonucleotide library against 23S rRNA investigate the structural effects of the probe hybridization on assembly.

**Fig.1.**
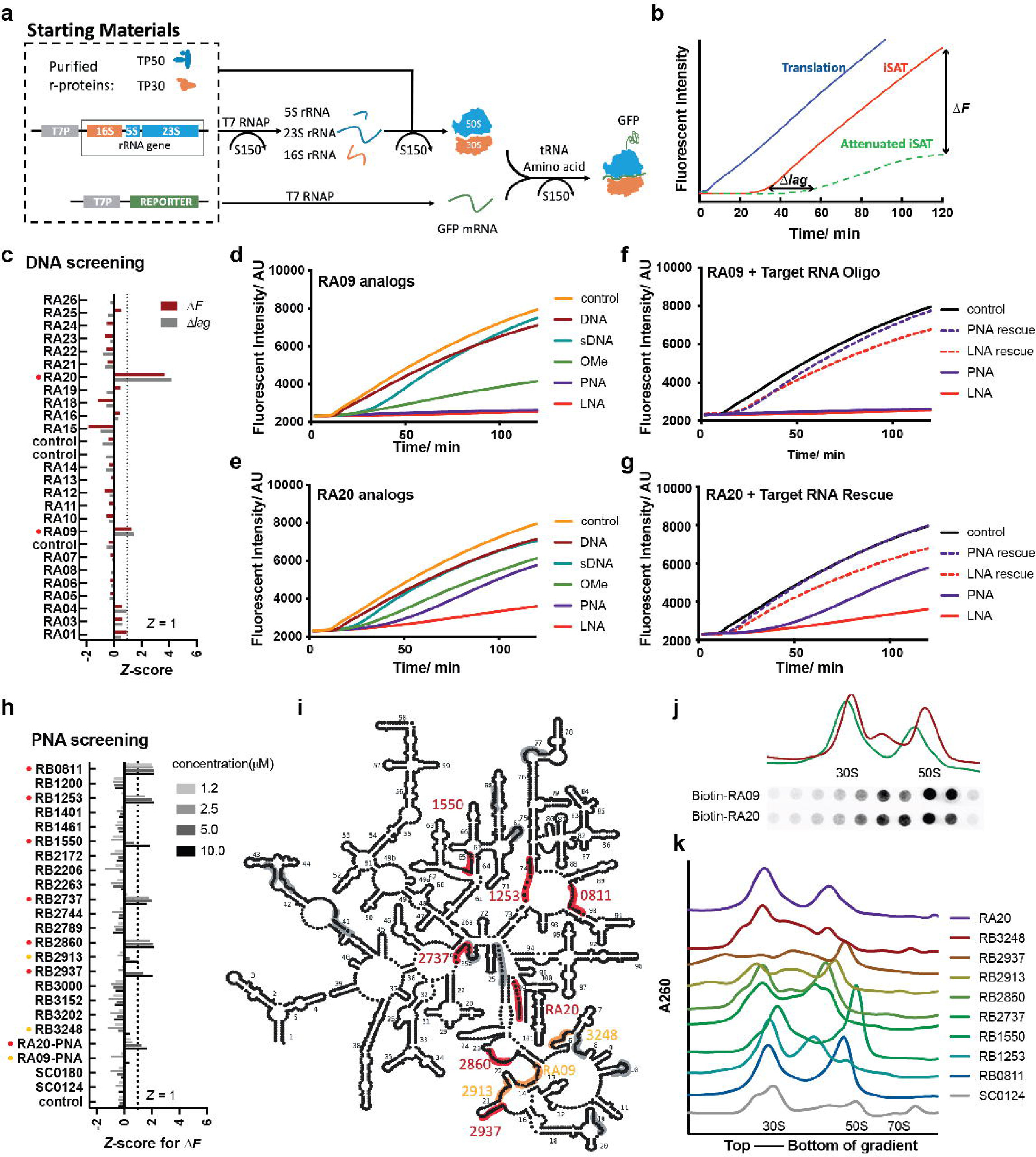
Attenuated iSAT assay for ASO assembly inhibitor screening. **a)** iSAT platform is performed in purified cell lysate S150, the rRNA will be transcribed and assembly with purified r-proteins *in situ*. **b**) This platform gives a window between in vitro translation and iSAT, enabled the possibility to monitor the time of lag (Δ*lag*) and fluorescent change (Δ*F*) for attenuated iSAT. In the first screening on ssDNA library **c**), two potential hits (red dots, RA20 and RA09) was identified with Z-score of either Δ*lag* or Δ*F* greater than 1. Their sDNA (cyan), 2’-OMe (green), PNA (purple) and LNA (red) analogs were tested and iSAT fluorescent curved shown in **d**) and **e**) respectively. The iSAT curve of rescuing PNA and LNA (purple and red dashed line) by antidote RNAs is shown for **f**) RA09 and **g**) RA20. **h**) shows the Z-core of Δ*F* for PNA screening for four different concentration (1.2, 2.5, 5.0 and 10 µM from lighter grey to black). The Strong attenuators with at least one Z-score is greater 1 are labeled with red dots, and moderate attenuators with at least one Z-score is greater 0 are labeled with orange dots. The strong and moderate attenuators are labeled on 23S/5S rRNA roadkill map in **i**). Sucrose gradient profiles for picked PNA from i) along with a control sequence SC0124 are shown in figure **j**). The 30S, 50S and 70S position are marked for the SC0124 trace. **k**) Streptavidin-Cy5 dotblot against Biotinylated PNA in sucrose gradient fractions for iSAT reaction treated with 1 µM Biotin-RA09 and Biotin-RA20. The dotblot was aligned with the 30S and 50S region with corresponding sucrose gradient traces.

It was convenient to use ssDNA probes for initial screening using ssDNA with a small set of 24 probes targeting regions distributed around the 16S and 23S rRNA secondary structure, and two potential hits were obtained that reduced ribosome production efficiency, and in some cases introduced an additional assembly delay. These initial hits were promising, but we explored use of several well-known analogs developed for increased hybridization stability for antisense and other biophysical studies, including 2’-OMe-RNA (OMe), peptide-nucleic acid (PNA) and locked nucleic acid (LNA). In general, these modified probes showed dose responses that were consistent with their well characterized *in vitro* hybridization energies as LNA > PNA > OMe > DNA. PNAs were chosen as compromise between stability and synthesis cost, and a 20-membered PNA library was screened in the iSAT reaction, providing 8 additional hits with high or moderate inhibitory effect. Intermediates that accumulate in the presence of 10 µM of 6 chosen PNAs were enriched using sucrose density gradient ultracentrifugation. Intermediate containing fractions were then analyzed using single particle cryo-EM, in the same manner as our previous reports for a variety of perturbations.^6,7^ In the 6 datasets, we identified a total of 47 pre-50S intermediates using iterative sub-classification, and of these, 39 intermediates were distinct from the ∼150 intermediates previously reported. Analysis of this extensive set of trapped intermediates, allows a much finer dissection of the 23S assembly pathway which was previously impossible. In particular, by trapping 50S assembly at an early stage, we are able to develop a kinetic model for intra-domain folding and inter-domain docking for Domain I, II, III and VI. Also, detailed step-wise assembly order for H32-H35 assembly first time showed structure evidence for templated-directed RNA foldon dockings, providing a possible general rule for efficient RNP assembly involving complete tertiary structures.

The inhibition of assembly by sequence-specific hybridization has potentially broader implications beyond providing key mechanistic insights on ribosome assembly. The need for new antibiotic development persists, and antisense oligonucleotides offer precise targeting advantages. While some studies have explored antisense targeting of essential mRNAs, direct inhibition of translation by targeting the ribosome is also under investigation.^20–22^ Our screening results and subsequent structural characterization highlight the potential of novel antibiotics targeting ribosome biogenesis, which could provide a productive avenue addressing a significant challenge in antibiotic development.

## Result

### Anti-sense oligonucleotide analogs block 50S-ribosome assembly

In the iSAT rection, there are two potential readouts of assembly inhibition, which are an elongated assembly window (Δ*lag*) and/or a lower final fluorescence (Δ*F*) at endpoint of the assay due to reduced production of functional ribosomes (**Fig.1 b**). In the initial round of screening, 24 single stranded DNA anti-sense oligonucleotides (ASOs) were chosen, with varied length and different target sites distributed across both 16S and 23S rRNA. Among all the DNA ASOs, RA20 and RA09 showed potential for impacting the ribosome assembly, as judged by that a *Z*-score over 1 for Δ*lag* (**Fig.1 c**) (See Methods). However, compared to control iSAT curve, the effect of DNA ASOs was subtle (**Fig.1 d,e**) and no significant changes were observed in sucrose gradient profile (Data not shown).

Reasoning that poor thermodynamic stability of the short DNA-RNA hybrids might limit assembly, ASO analogs harboring different backbones were examined. OMe, PNA, and LNA analogs of RA20 and RA09 indeed showed improved inhibitory effects in the iSAT reaction, and PNAs and LNAs almost completely shut down the translation of GFP (**Fig.1 d,e**). To show specificity, a rescue experiment with single stranded RNA antidotes were performed, the pre-hybridization of PNA/LNA to the complementary antidote RNA significantly restored the efficiency of the iSAT reaction (**Fig.1 f,g**). A scrambled sequence of RA20-PNA was used as a control for sequence-specific hybridization, and indeed the scrambled probe showed no iSAT inhibition (**Extended Data Fig.1 b**). Further, the sucrose gradient profile of the PNA treated reactions indeed showed accumulation of intermediates due to the targeted inhibition of assembly. The inhibition of the GFP reporter expression is concentration dependent, accompanied by increased accumulation of pre-50S on sucrose gradient trace (**Extended Data Fig.1 a-b, e-f**). To provide a potential affinity handle for trapped intermediates, RA20-PNA and RA09-PNA were resynthesized with a biotin-tag, and both biotin-RA20-PNA and biotin-RA09-PNA showed a similar inhibitory effect, and the biotin tag was colocalized with the intermediate peaks in dot blot against streptavidin-Cy5 conjugate (**Fig.1 j**)

### Secondary Screening with an expanded PNA library

Having demonstrated PNAs as more potent iSAT inhibitors compared to the initial DNA probes, a second round of screening was performed with, twenty additional 13-mer PNAs were selected and synthesized and tested in iSAT assay (**Fig.1 i**). In this set of experiments, it was difficult to accurately quantify the delayed window (Δ*lag*) due to the flat nature of many of the curves. Thus, end-point GFP fluorescence with varied probe concentrations was used to assess the inhibitory effect of the second library. Half of the PNAs exhibited a clear dose-dependent inhibition as judged by the end-point GFP fluorescence (**Extended Data Fig.1 e**). Of these, 9 PNAs with a Z-score greater than 0 to move forward. 5 out of 9 has a Z-score greater than 1. (**Fig.1 h**) Again, altered sucrose gradient profile showed up with feature of decreasing 70S/polysome and emerging of pre-50S peaks (**Fig.1 k**). The differential sedimentation of pre-50S implies different targeting site resulted in varied assembly intermediates with distinguishable hydrodynamic radii. Quantitative proteomics of sucrose density gradient fractions demonstrated the compositional heterogeneity resulting from different PNA oligos. (**Extended Data Fig.2 a**) For example, RB2937 showed depletion of uL29 and RB2860 showed depletion of bL34. The r-proteins are located near the targeting site in both of these cases. Also, some intermediate peak in PNA-inhibited dataset showed enrichment of LSU assembly factors such as YjgA and YggL were observed in the intermediate fractions. (**Extended Data Fig.2 b**) Yjga was hard to resolve in iSAT dataset. However, intermediate fraction in RB1550 showed an increase in YjgA levels. It targeted H65, an element within the uL2 block in previous assembly dependency map. In bL17-depletion dataset, the attachment of YjgA in the D or E class exhibited mutual exclusivity with the uL2 block. This implied the RB1550 possibly disrupted the H65 and thus the YjgA binding site was more exposed.

### 47 new pre-50S electron density maps showed PNA specificity and detailed rRNA assembly dependency

Pre-50S and 50S fractions for six PNA ASOs along with control iSAT reaction were pooled and intermediate density maps were solved using CryoEM using our previously described workflow with slight modifications described in Methods.^7^ Overall, 47 new pre-50S density maps were reconstructed. To examine structure variation, we performed hierarchical analysis that pairwise compared with previous classes^3,5,7^ (**Extended Data Fig.3**). All the intermediate density maps are organized into a maturation order in **Fig.2 a**. As previously defined, B classes are early classes that have majority domain I, II, III an VI assembled. The G classes lack formation of domain II, while the C classes are based on B classes, with additional maturation of the base, and the E classes are the most mature class preceding a complete 50S subunit.

**Fig.2.**
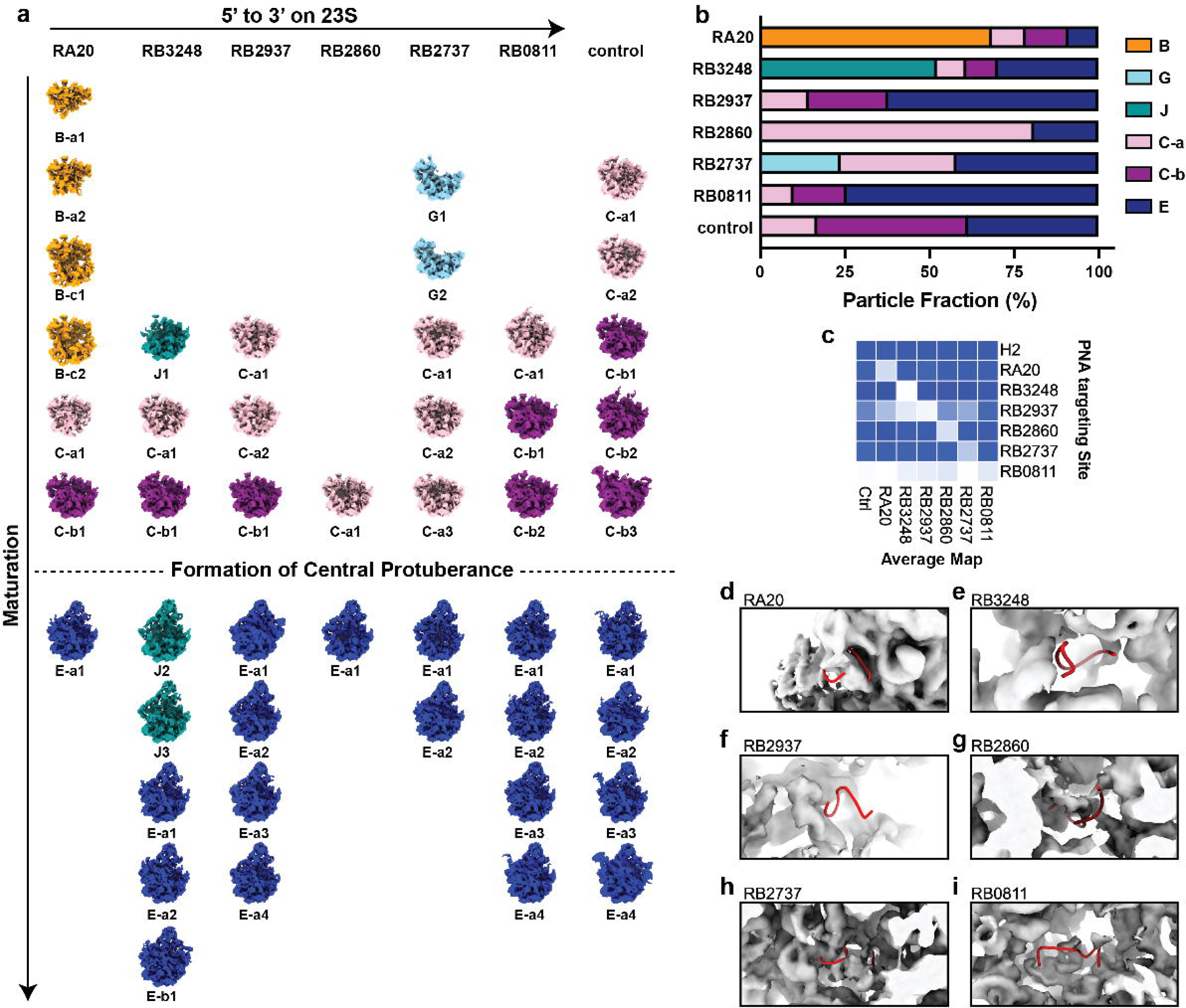
47 density maps for attenuated iSAT intermediates. **a)** intermediates layout for 47 arranged for different PNA treated samples (columns ordered by targeting site from 5’ to 3’. On the rightmost, iSAT without treatment serves as a control sample). Rows are ordered by maturation status from smallest to largest for each dataset. The intermediates were aligned in the center. The classes for maps are labeled below each thumbnail and colored accordingly (B: orange, G: light blue, J: cyan, C-a: pink, C-b: purple and E: darkblue). **b**) The class fractions for different samples are shown in barplot. c) The mean electron density in targeting site mask is calculated for weighted average map for each class in **c**). The y-axis shows targeting site for different PNAs. H2 serves as a negative control. The x-axis showed name for average maps. The blueness is proportional to the mean electron density. **d-i**) shows the PNA targeting site (red ribbons) on corresponding average map (from **d** to **i**: RA20, RB3248, RB2937, RB2860, RB2737 and RB0811). The control average map was in Figure S4.4.

In addition to these classes previously observed classes, a new major class appeared in the hierarchical analysis **(Fig.2 a**, **Extended Data Fig.3**). In the presence of RB3248, there are three intermediates, now named as the J class, which have major defect in the assembly of domain I helices. Also, in RA20, we observed a new subclass B-c, where the majority of Domain I and II is formed and in addition, the majority of domains III, and VI are formed, but the docking of these two regions is hindered, leading to a large non-native assembly intermediate. The vast majority of previously observed intermediates have exhibited largely native-structure, few such non-native arrangements.

Different PNA-inhibited datasets showed various particle distributions among the major observed classes (**Fig.2 b**), which aligned with variations in sucrose gradient profile (**Fig.1 k**). In general, the closer PNA targeting site to 5’ end, there are higher probability to resolve earlier assembly intermediate. RA20, which targeted the most 5’ helix (H1) in 23S produced the earliest B class. RB2737 which targeting H102/H26, the junction of Domain II, led to accumulation of G class which does not have assembled domain II. On the other hand, RB0811 which binds the PTC region near the 3’ ends on 23S, have the most abundant E class.

Interestingly, when we scrutinized the targeting site in different PNA-inhibited datasets, in most of the cases, the electron density surrounding the targeting site was conspicuously missing (**Fig2. c**), which is consistent with the idea that hybridization of the PNA disrupted the native folding and made the region more unstructured or too flexible to be resolved using CryoEM (**Fig.2 d-i**). In the control sample, aside from the late assembly helix targeted by RA0811, the electron density surrounding the same targeting site appears normal. (**Extended Data Fig.4**)

### 3D flex reconstruction facilitated the structure solving of RA-B-c class

Drastic conformational changes were observed in H1 folding. Comparing RA20-B-c1 and RA20-B-c2, the major difference is the movement of the domain III and VI. Although the conformation must be sufficiently stable to be captured by cryoEM, domain I/II were primarily still driving the alignment of particles, and the bottom density for domains III/VI were not well-resolved in the initial heterogeneous subclassification. Analysis of this subclass was facilitated by application of the 3D Flex model in CryoSPARC^23^ to capture the movement. With the trained model in 3D Flex, we are able to have a more reliable 3D Flex reconstruction on the RA-B-c class after combined particles from RA20-B-c1 and RA20-B-c2. (**Extended Data Fig.5**) Compared with homogeneous refinement, the Domain I/II regions were similar, while the 3D Flex showed clearer features of RNA helix feature for domain III and VI.

### Comprehensive Domain I/II/III/VI rRNA assembly for 50S

In order to get a detailed helix assembly hierarchy, we manually curated the RNA structure elements using defined helical elements and proteins as structure components (**Supplementary Material 1**). Using our previous developed occupancy analysis, we assigned formation of structure elements for each intermediate by thresholding the normalized electron density occupancy. (**Extended Data Fig.6 ab**). The occupancy matrix was then used to develop a contact dependency using two criteria. First, information about the dependency of two elements is revealed in a scatter plot of their occupancies. Dependencies are revealed by the clustering of points in particular quadrants (Q1-Q4), indicating one element never forms without the other one being pre-formed. For example, in **Fig.3 a**, the occurrence of H103 is after bL34. Second, these two elements need to form native contacts in the mature 50S **(Fig.3 b**). Using network analysis, a comprehensive contact dependency map for Domain I/II/III/VI rRNA assembly was composed. All the dependency is listed in supplementary data. In order to further simplify the graph, the dependency was pruned redundant edges using previous methods and only elements in domain I, II, III and VI are considered to focus on early assembly events. Finally, a comprehensive contact dependency graph was generated. (**Fig.3 c**) Pairs that only have Q1 and Q4 dots are cooperative structure elements. They will be grouped into assembly blocks and shown in shaded boxes. For example, uL29 and H6 assemble cooperatively.

**Fig.3.**
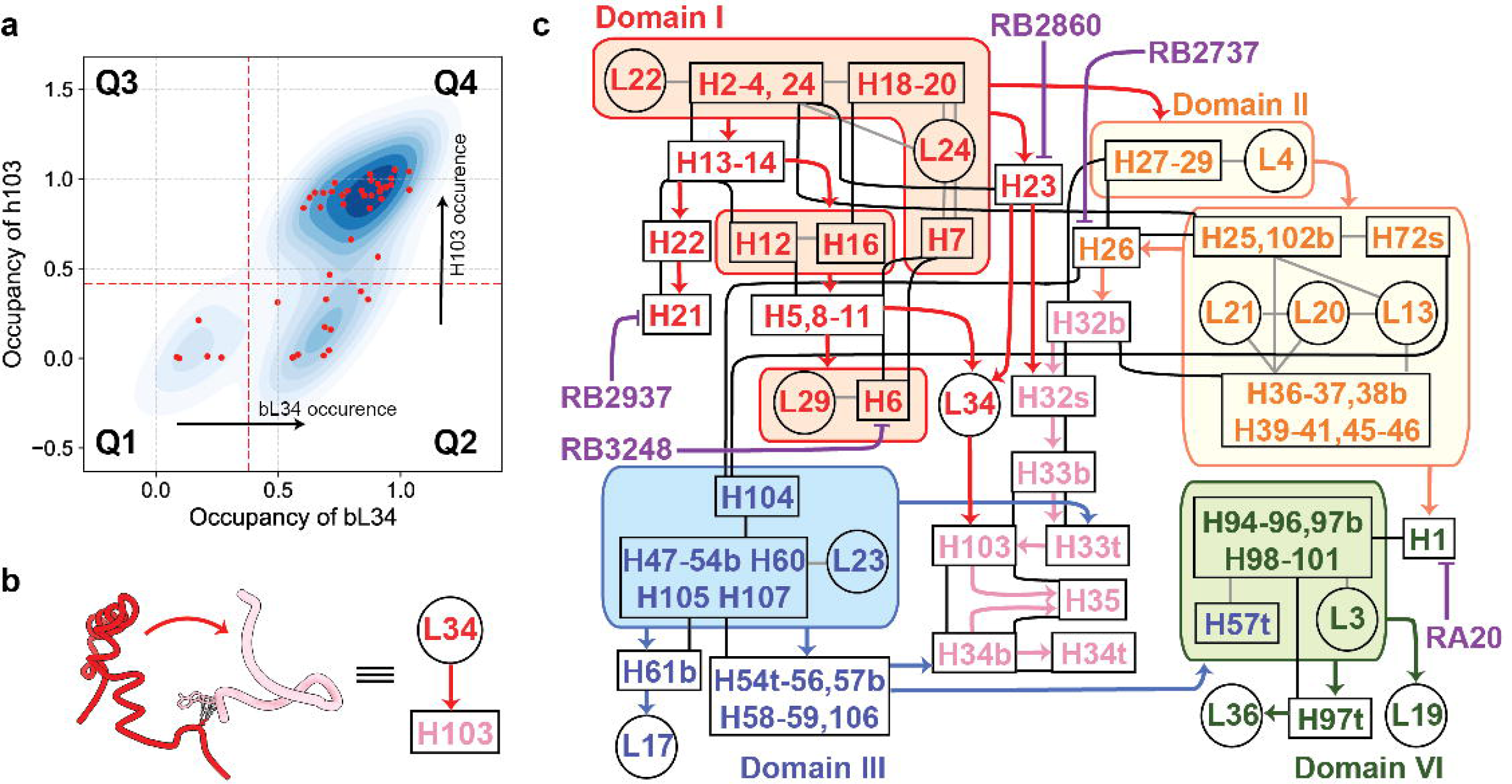
Comprehensive contact dependency graph for 50S domain I, II, III and VI. **a)** Quadrant scattering kernel density plot for bL34-H103 mean electron density, in which each black dot represents an intermediate map. Dots in Q1 and Q4 indicates that bL34 and H103 are both missing and assembled respectively. Q2 implies bL34 occurrence happens before H103. **b**) grey dash line shows native interaction between bL34 and H103, together with the occurrence order in a), the contact dependency from bL34 to H103 was drawn. **c**) A contact dependency graph for domain I, II, III and VI. Cooperative structure elements are grouped in shaded boxes, the tertiary contact or protein-RNA interaction in assembly blocks are shown in grey lines. The covalent linkages between helices are shown by black lines. Contact dependency was bold arrows colored by domains of the parental node (red: domain I, orange: domain II without H32-H35, H32-H35: pink, domain III: blue and domain VI: green.). Disruptions of helix formation by different PNAs are shown in purple blunt arrows.

## Discussion

### Intra-domain folding is independent from inter-domain docking

In the RA20-B-c class, the H1 is targeted with PNA-RA20. The most 5’ end and 3’ end of 23S form the H1, which connects domain I and domain VI 23S rRNA in a close loop. The hybridization of H1 is also a milestone for LSU assembly process subsequently followed by processing by several nucleases.^24^ In a matured LSU, H1 contacts with domain VI through H94 and H98 and stabilize the LSU.^25^ From previous observations, compared to RA20-B-a1, in deaD-B-a1, the H1 formation was observed without other parts in domain VI formed (**Extended Data Fig.7 ab**). Also, it is clear that the docking of domain III does not need domain I with observation of deaD-preB2 (**Extended Data Fig.7 bc**). In RA20-B-c, the density of H1 was also dampened like other RA20 maps, but interestingly, the both domain III and VI were folded, but with a drastic conformational change.

This is the first time a large-scale non-native domain arrangement was observed in LSU assembly process in bacteria, which provided a new insight into LSU assembly. In the model fitted in RA20-B-c, with H1 as a broken hinge, the majority part of domain III and domain VI are locked halfway during the docking process, where Domain I/II/III/VI has folded but the consolidation of Domain I/II and domain III/VI to get a C-a class got stuck (**Fig.4 a-c**). This open conformation resembles a lazy-8 from the solvent side view (**Fig.4 d**), with one circle on the left surrounded by domain II and mis-docked domain VI, and another circle is the opened exit tunnel. Three anchor points between domain I, II (the upper part) and domain III, VI (the lower part) are mediated by three r-proteins, D-II-uL13-H98, D-I-uL22-H50, and H51-uL23-uL29/H9 (**Fig.4 d**). These r-proteins help to glue the assembled upper and lower part and stabilize the open conformation. Notably, loops in H50 in domain III and H98 in domain VI adopted a non-native conformation in order to keep contact with uL22 and uL13 respectively. The 5’ half of H104, which the only covalent linkage between domain II and domain III, is stretched out without any other specific interaction. According to 3D Flex generated movies, (**Movie 1, 2**) the domain III/VI can still move around, interestingly, these anchor points are always remained, which implies the interactions are to some degree, stable and specific.

**Fig.4.**
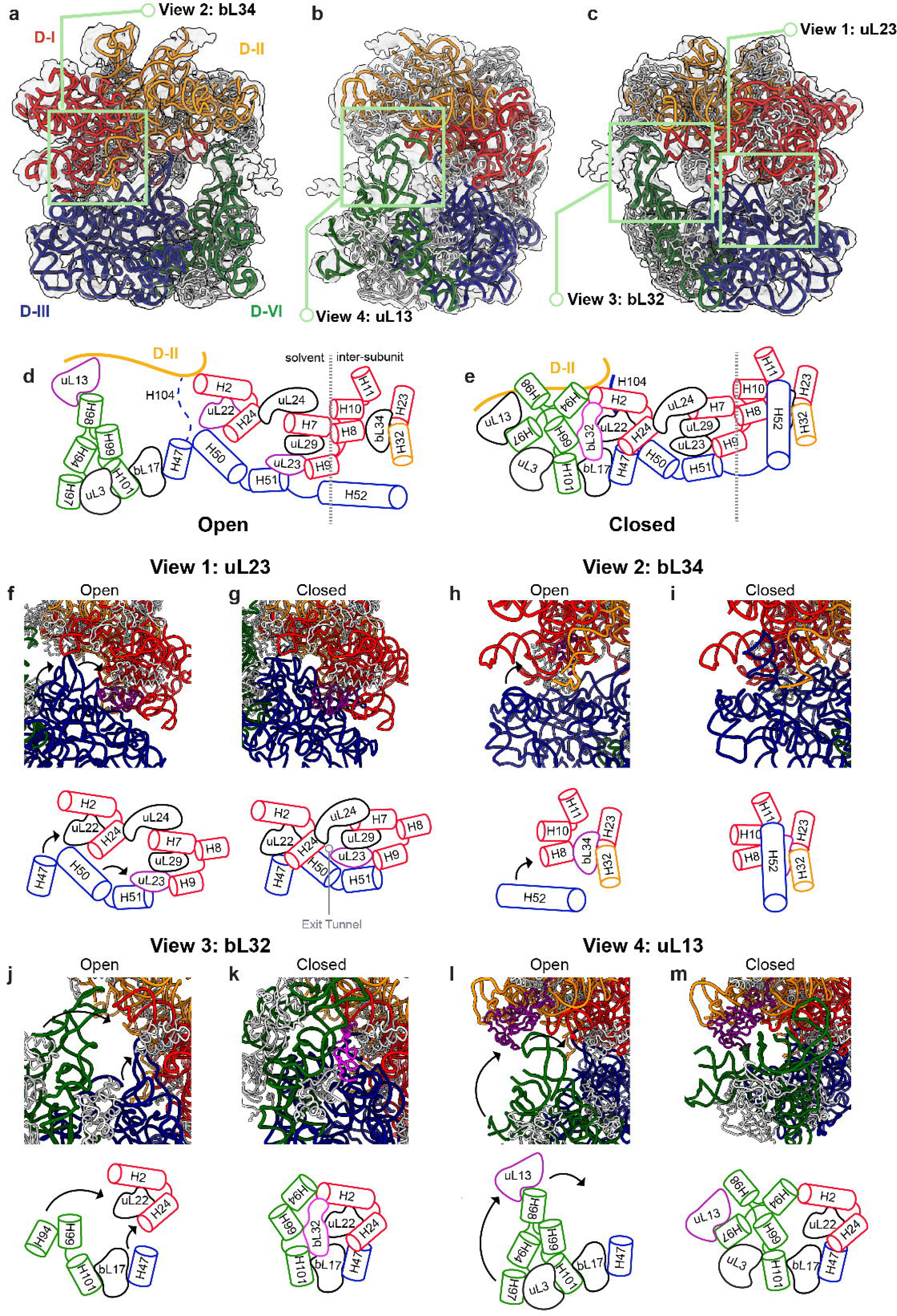
Drastic large-scale non-native domain arrangements in RA-B-c class. **a)** inter-subunit, **b**) side and **c**) solvent-side views for RA-B-c atomic model shown in electron density map (transparent grey). The 23S RNA are colored according to domains (red: domain I, orange: domain II, domain III: blue and domain VI: green.) The ribosomal proteins are colored in light grey. Simplified models for key components in domain III and VI docking were shown for **d**) open conformation and **e**) close(native) conformation (PDB: 4YBB). RNA helices are displayed by cylinders using the same colors in **a-c**). Most proteins are colored in black. Three important anchor proteins are highlighted in purple in **d**). bL32 was shown in pink in **e**). Simplified models are flattened, and the dashed line in d and e indicates boundary between solvent and inter-subunit views. **f-m**) zoomed in views for green boxes in **a-c**) centered in highlighted proteins with corresponding simplified models below. **f**) uL23 view, **h**) bL34 view, **j**) bL32 view and **l**) uL13 view. **g, i, k** and **m** are corresponding close conformation. The centered protein is colored in purple or pink. Black arrows in open conformation demonstrate the motion from open to close conformation.

A movie illustrating the morphology change between this open conformation and closed (native) conformation was generated (see Methods) (**Movie 3**). Based on that, a simplified model with only showing the key components involving in the docking process is demonstrated in **Fig.4 d-e**, which provided a possible cooperative docking mechanism. First, from uL23 view (**Fig.4 f-g**), uL23 adapted a large conformational change to glue uL29 and H51 together, which serves has an initial anchor point during the docking. The H47 interaction with uL22 and H24 replaces the non-native interaction between H50 and uL22. These two events mark the closing of the solvent side exit tunnel and success docking between domain I and III. From bL34 view (**Fig.4 h-i**), the bL34 sits in the native cleft formed between domain I and domain II. The docking drives the lid closure of bL34 binding site with H52, revealing the mystery of bL34 cavity formation that is deep inside of the 50S after maturation. Also, H52 extensively interact with H8-10 and bL34, with bL34 interaction with domain II rRNA through H32. Domain I, II and III interlocked at bL34 side and provided further stabilization. bL32 is not present in the open conformation (**Fig.4 j**). In the Nierhaus map and previous bL17 depletion result^3,26^, bL32 showed strong dependency on bL17. However, in the open conformation, bL17 was already incorporated but bL32 was not. The depletion of bL32 was also confirmed by proteomic data for intermediate fraction (**Extended Data Fig.2 a**). This implied the binding event of bL32 does not solely depend on bL17 but also needs the correct formation of junctions between domain I, III and VI. bL32 also extensively interact with these three domains and possibly provide further stabilization (**Fig.4 k**). The non-native interactions between helix and proteins are firstly reported during the 50S biogenesis (**Fig.4 l**). This interaction between H98-uL13 closed the left circle in the 8, and therefore we were able to capture this intermediate conformation. This non-native interaction is also possibly happening in during the normal assembly process, that will presumably help with the directing and positioning of domain VI rRNA. During maturation, the native H97-uL13 will then replace the non-native interaction (**Fig.4 m**). All of these dockings did not need domain II participation, which supports the basis of G-class observations.

In general, even in a matured C-class where upper and lower parts get docked correctly, there are a lot of contacts between RNA helices among these domains. However, in the open conformation, the helices between the upper and lower part have no contact, which suggest the folding of domain III and VI does not require a pocket or a template from domain I and II. In the other words, the intra-domain folding of rRNA is independent from inter-domain docking, and instead, the protein connections are more essential for specifically guiding the joint of these two parts. The independent domain folding also aligns with the thought of parallel assembly of LSU which is one of the most efficient ways to make a ribosome.

### Intra-domain folding is earlier than inter-domain docking

Recent studies on assembly of circular permutated RNA showed that the docking hierarchy resembled the normal order of 23S rRNA.^27^ In one of the circularly permuted constructs starting from H45, which is the end of domain II rRNA, no intermediates where domain III/VI were ordered were observed. The present observation of independent folding of domain III and VI rRNA, implies that the folding of intra-domain can occur earlier than inter-domain docking, even though the undocked domains III and VI were not previously observed. There is a possibility that the intradomain folding is faster than the docking, so that any folded domain III and VI are quickly docked with domain I and II. The intra-domain folding steps are more likely to be a rate-determining step compared to docking of correctly folding blocks that possibly has already oriented by primary constrains.

### Templated-directed RNA foldon docking and domain consolidation

Despite the intra-domain folding and inter-domain docking mechanism of the majority of the core structure constituting the solvent side of the LSU, then later part of rRNA folding follows well organized pathway mediated by tertiary interaction, protein scaffolding with rRNA primary sequence connection constrains, which is by PNA probing.

For example, in middle part of the contact dependency map, the smallest structural elements for H32-35 assembly is primarily impacted by two PNAs, RB2737 and RB2860. Compared with other PNA-inhibited datasets, there are no C-b class for both RB2737 and RB2860. An extreme case in RB2860 is all its intermediates arrested in one of two states, one C-a class, RB2860-C-a1 and one E-a class, RB2860-E-a1, the primary difference being RB2860-E-a1 has mature central protuberance. They both are lacking H32-35 and the whole uL2 block. This result aligns with the previous dependency map very well, where without formation of H32-35, the H63 and the uL2 block cannot correctly dock, while the central protuberance formation does not require H32-35.

These two datasets demonstrated a detailed pathway, at single helix resolution, of how RNA foldons in H32-35 dock onto a previous folded pocket, along with the consolidation of the scaffold of domain I, II and III 23S rRNA. In our observations, the smallest RNA foldons are often hairpin loops, designated as H[number]t, t for tip. For most cases, as H32-35 exemplified, when the RNA has secondary structure where a double helix is formed, the foldons will be the intact double stranded RNAs. We did not observe any independent folding of a split single stranded RNA which is present in a helix in the matured 50S. (**Fig.5 a**)

**Fig.5.**
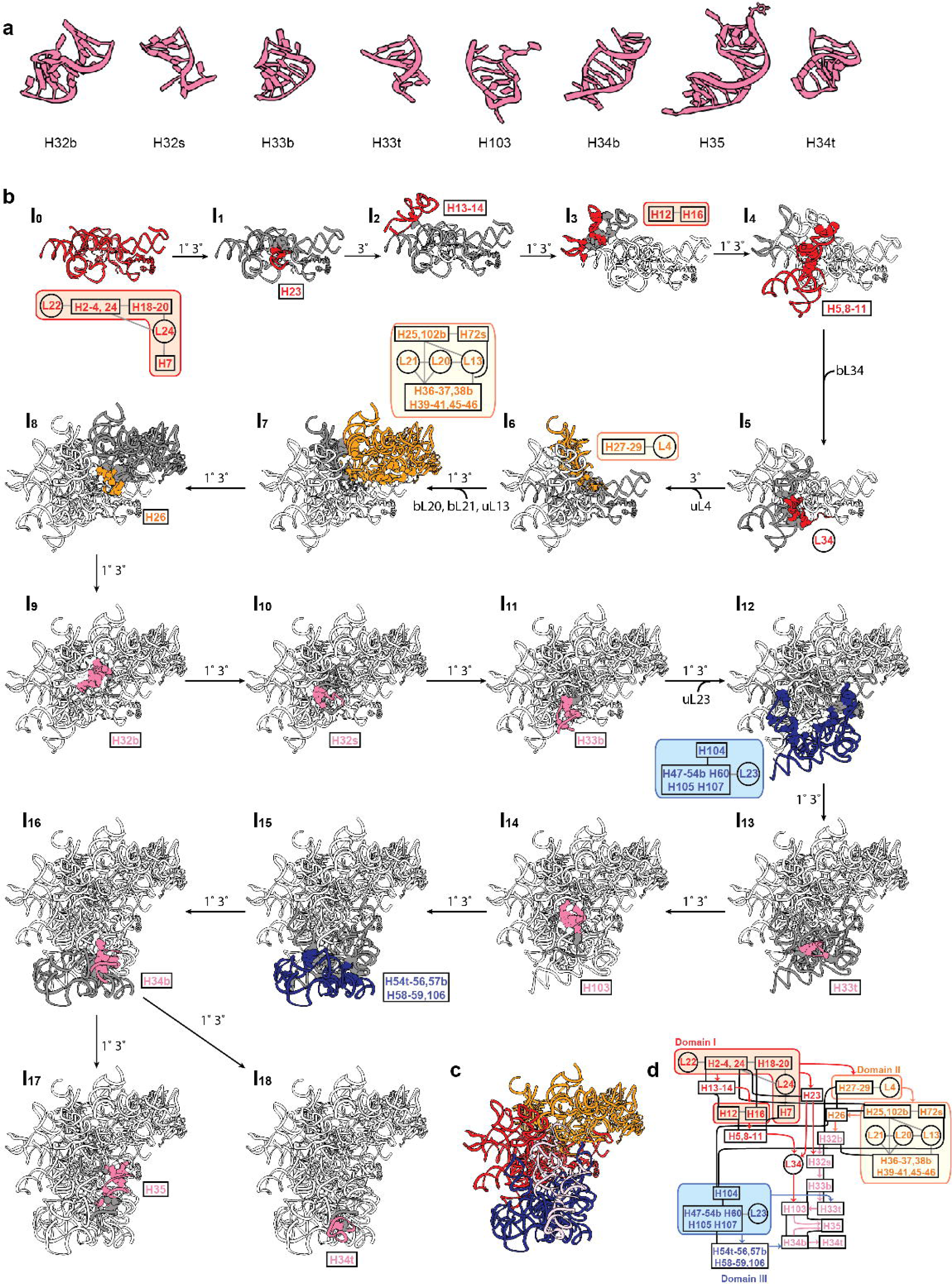
Templated directed foldon docking and domain consolidation during H32-35 assembly. **a)** RNA foldons in H32-35, ordered by occurrence. **b**) Assembly pathway of H32-35. The newly assembled structure was colored according to its domain with its content from contact dependency graph displayed close to it (red: domain I, orange: domain II without H32-H35, H32-H35: pink, domain III: blue and domain VI: green). The parental node is colored in dark grey. The interactions between newly assembled structure and previous steps are shown in balls. From I_0_ to I_16_ is a linear pathway for H32-35 assembly, each step is forming the minimal requirement for latter step of H32-35 assembly. From I_16_ to I_17_ or I_18_ are independent assembly for H35 and H34t, respectively. 1° and 3° along the assembly order indicates covalent connection in RNA and tertiary interaction between the newly assembled structure and the previous intermediate. **c**) shows over all assembled atomic models for this pathway with their connections from contact dependency graph in **Fig.3 c**. All atomic models are truncations of PDB:4YBB

The pathway (**Fig.5 b**) from I_0_ to I_7_ shows the minimal requirement for starting assembly the H32-35. H26 in I_8_ is the important junction in domain II, which has extensive tertiary interaction with both domain I and II rRNA. It also has primary connection with H32b, the starting point of assembly the H32-35 block. From I_9_ to I_11_, the H32-H33b begins to fold through both a primary connection with preceding sequences, also into the scaffold made by domain I and II through tertiary interactions. In order to correctly dock the remaining helices in the H32-35 block, the domain III rRNA needs to fold first. For example, H33t needs the docking site on uL23 block (I_12_ to I_13_), and H34b needs scaffold of H54t-H59 (I_15_ to I _16_). Overall, each step in folding of H32-35 is well sequenced, in that each section has a sequence connection with regions formed in the previous step, which constrains the distance of the RNA foldons before they dock into their now-formed pocket on the scaffold via tertiary interactions and/or protein-RNA interactions.

### Conserved smallest scaffold for ET_solvent_ formation

In this dataset, we included three domain I targeting PNA RB3248, RB2937 and RB2860. RB3248 targets the H6, which immediate blocks the uL29 binding. It has also impacted the assembly of H5,8-16, significantly in all three J class in RB3248 dataset. In RB2937 dataset, H20-21 are missing, and at the same time all intermediates have a common defect on L1 stalk by destroy the anchoring site for L1 stalk base, H75 and H79. In RB2860, the density for H26 and H26 binding protein, bL34 were both missing.

With consolidated evidence in the domain I-targeted PNA-inhibited dataset, we are able to get the smallest consensus structure element for all the intermediate maps. The smallest scaffold is shown in the upper box of dependency plot, which contains domain I H2-4,24, H18-20, H7, and two r-proteins, uL22 and uL24. (**Fig.6**) Surprisingly, the RNA helices in this small block are solely bridged by proteins with only a few inter-helix contacts. uL24 plays an important role of clamping these three helices together and uL22 holds the H2-4,24 together and also interacts with all domains after domain I. (**Extended Data Fig.8 ab**).

**Fig.6.**
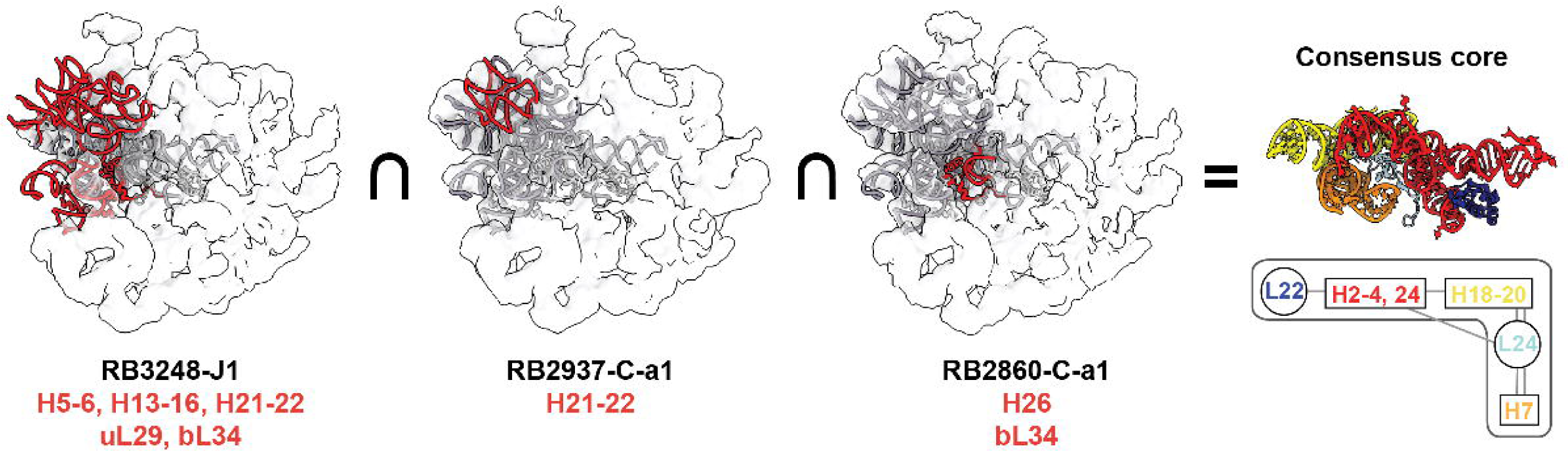
Consensus core obtained by intermediate intersection. Smallest intermediates density maps from three domain I targeted PNA dataset (from left to right: RB3248-J1, RB2937-C-a1 and RB2860-C-a1) are overlaid with 4YBB model of uL22, uL24, bL34 and domain I rRNA. The atomic model of missing parts in the intermediate are colored red, while the existed parts are colored grey. The intersection of the three models is shown on the rightmost with the simplified representation in the contact dependency map in **Fig.3 c**. (uL22: dark blue, uL24: light blue, H2-4,24: red, H7: orange, H18-20: yellow)

In a previous study^7^, we showed that the ET_solvent_ side formation serves as a scaffold of LSU formation. Without PNAs, it was not possible to dissect the 600 nucleotide and 4 proteins into smaller blocks. However, using PNAs, we are able to further dissect the domain I into smaller pieces, leaving the most important part for building up the upper part of ET_solvent_ scaffold. Interestingly, in RB3248 dataset, it created a large missing piece in H5-6, H8-14 which is the structure ortholog of 5.8S in higher organism. (**Extended Data Fig.8 c-e**)

The sequence of the core scaffold of rRNA sequences are not highly conserved among species, however, the structure of the scaffold is highly conserved with the clamping of uL24 and uL22 among all domains of life. **(Extended Data Fig.9 a**) In some small genome, such as microsporidians and yeast mitochondria, different degrees of truncations in H7, H18-20 and H24 were observed. (**Extended Data Fig.9 b**) In yeast mitoribosome, the uL22, uL23 and uL24 make new close contacts. This maybe a possible way to compensate the truncations of H24. Similarly, the uL24 also occupied the original position in H7 and makes contact with other helices and r-proteins with largely extended C-terminal.

In a remarkable deviation, human mitochondrial ribosomes exhibit extreme truncations in H24, H18-20, and H7. (**Extended Data Fig.9 c**) Additionally, most helices in the exit tunnel domain I are replaced with mitochondrial elaborated or supplementary r-proteins from host genome. For example, the uL22 and uL24 are both extended, and the original H24 was replaced with mL45, which could be due to mitochondrial gene compression, exit tunnel specialization, and an alternative pathway for this region in ribosome assembly.

### Platform for study of a large RNP complex

There are multiple rDNA operons in most bacteria, 7 in *E. coli* specifically, which made it difficult to perform genetic manipulation directly on rRNA. Even with the “Δ*7*” strain in which a deletion of all 7 operons was complemented with plasmid encoding one of the operons, mutation of rRNA sequences is tricky and extremely unfeasible for lethal mutations. iSAT instead, provides a convenient and clean platform for screening lethal mutations and ribosome assembly inhibitors.

Using bacterial LSU as a model, we are able to utilize the complementary oligonucleotide analogs to site specifically probe the folding of large RNP complex. With the cases we’ve shown, the probes are able to compete with native RNA helix formation, RNA-RNA tertiary interactions and RNA-protein interaction. These perturbations can easily result in corresponding RNP assembly intermediates for further structural characterization which in our study with single particle cryoEM and proteomics and can be easily expanded to existing sequencing-based RNA structure probing. Single molecule fluorescence has been crucial in studying how ribosomal proteins bind over time, revealing how the ribosome assembles during transcription^28–30^. By using fluorescent tagged ASOs, we can delve deeper into this kinetics, offering a clearer picture of RNA folding in a co-transcriptional assembly model.

Different backbones and concentration should be considered in order to get optimized specificity. We observed a consistent order of inhibition with different ASO analog backbones in RA09 and RA20, which is:

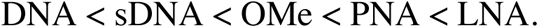

This series is consistent with melting temperature of duplex of these oligos with complementary RNA (**Extended Data Fig.10 a**), with an exception of Tm of sDNA/RNA lower than DNA/RNA. This could be a result of nuclease resistance of sDNA compared to DNA. For the RA20-LNA, we also tried different locked site for the oligo (**Extended Data Fig.10 b**). It demonstrated different locked site will have impacts on inhibiting the RNP assembly, probably resulted from varied seeding site of the binding. Also, the combination of 2’ lock and phosphothioate backbones have additive impact also suggesting there are other factors for increasing kinetic blocking for sDNA since 2’ lock already has nuclease resistance.

While most ASOs showed and inhibitory effect, we also observed a chaperone effect from several ASOs. For DNA-RA15, it also showed shorter lag time and higher fluorescent signal, implying that assembly efficiency was increased. Further, in the PNA-RB0811 dataset, there are more E class particles compare to control iSAT reaction. RB0811 binds to highly modified PTC, which might mimic the effect of assembly factors. Also, DNA-RA15 binds to the domain II junction, where is the potential functional site for DEAD-box helicases SrmB and DeaD. Both of these two ASOs might have similar roles of holding the unfolded transcript that helps to facilitate earlier assembly events.

The guiding principle of this work is that the ASOs are competing with native structure formation, operating in a kinetic window where the target is accessible prior to sequestration during the assembly process. Half of the PNAs showed no effect, and some of the PNAs showed only moderate inhibition, which could be either the assembly is slowed down and resolved afterwards, or the PNA failed to compete with the native structure formation. In those cases, the solved structures are mixture of PNA targeted and native intermediate. This information is useful for estimate kinetics. However, for characterization intermediate structure, one could perform potential pull-down and enrich the PNA bound structure.

### An early 50S assembly model from ASO perturbation

With different PNA binding, the overall of the assembly landscape is altered in three different scales that we have already discussed: a) Interdomain docking, b) details of RNA foldon and domain consolidation and c) non-essential parts in domain I scaffold for late assembly. All of these are summarized into our new early 50S assembly model with kinetic implications. (**Fig.7**)

**Fig.7.**
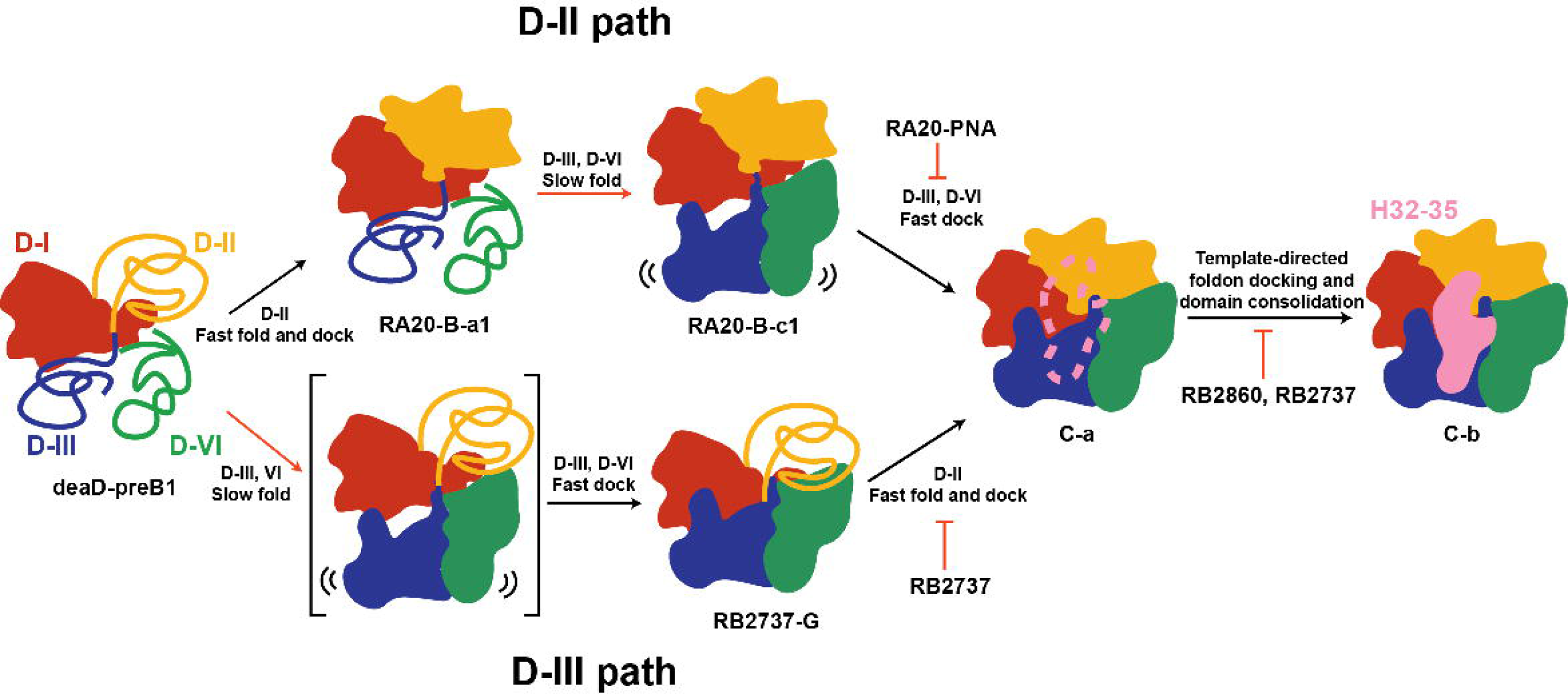
Kinetic assembly model for domain I, II, III and VI. A Scheme of overall kinetic assembly model from domain-I (D-I, same for II, III and VI) to C-b is shown. The domains are colored accordingly. Assembly of one domain is illustrated by solid blob while unfolded RNAs are shown in coils. There are two major pathways from preB1 to C-a, namely D-II (upper) and D-III (lower) path. The black arrows show faster steps while the red arrows show the slower steps. The PNA inhibition for different folding steps are shown in red blunt arrows.

In the early 50S assembly kinetic model, we are not able to catch a domain I-only intermediate in the PNA-inhibited datasets. But the domain I-only intermediate has been independently discovered in Δ*deaD* strain at low temperature and time-coursed *in vitro* reconstitution.^7,12^ After domain I rRNA is folded, there are two major pathways, D-II and D-III, in which domain II forms first or domain III forms first respectively. In RB3248, RB2937, RB2860, RB0811 and control datasets, it is hard to see an intermediate before C-a, likely because these processes are all really fast in the absence of perturbations. In the previous time course study of 50S assembly in iSAT, the B class (domain III, VI defect) and G class (domain II defect) were also observed, but only in early time points with a relatively small particle fraction.

In the D-II path, domain II forms first. RA20-PNA blocks the H1 formation, subsequently impacts the docking of domain III and VI. The ability to trap the B-class with PNAs is the basis for the proposed fast kinetic of Domain II folding and docking and slow kinetic of folding of D-III and VI. With coincident stabilization of anchor points described above, we are able to observe RA-B-c, which otherwise would not be easy to see because of fast docking kinetics of Domain III/VI to domain I/II.

In the D-III path, RB2737 blocks the H26 an important junction of domain II. It slows down the fast folding a docking of domain II, so that we can trap the G class after domain III, VI folding and docking. The fact that we are not able to capture the un-docked domain III and VI intermediate in D-III path, implies that the docking is relatively fast. Interestingly, G-classes are a common feature of intermediate in DEAD-box helicase strains, Δ*deaD* and Δ*srmB*, which could reflect a general or specific role for helicases in facilitating domain II folding and/or docking.

The 23s rRNA region termed Domain 0 was previously identified based on secondary and tertiary structure analysis.^31^ Domain 0 appears as a central hub, with an assumption of independent folding capacity. In this work, we can localize domain H26, H104 and H72s, H102b in sequence in our new contact dependency map (**Fig.3 c**). H26, H72s and H102b are important junctions for Domain II assembly. H104 connected domain II and domain III and is essential for Domain III docking. H104 and H26 form independently which aligns with D-II and D-III pathway respectively. In this work, domain 0 consolidation is not only a hallmark for ET_solvent_ maturation, but also marks the convergence to C-a class, followed by H32-35 formation to form a C-b class, which now is ready as a scaffold for domain VI and V folding and docking.

On a broader scale, our study elucidates the sequential and parallel docking order of ribosomal domains, providing insight into the overall assembly process. Furthermore, we offer detailed insights into the assembly principles governing RNA helices, particularly exemplified by the meticulous template-directed RNA foldon docking observed in the case of H32-35. This mechanism serves as a foundational principle driving the precise and controlled assembly of the 50S subunit, with potential applicability extending across structured RNA assembly kinetics.

## Methods

### Synthesis of Anti-sense oligonucleotides and their analogs

All DNA, sDNA, LNA, RNA and 2’-O-Me RNA were synthesis at Integrated DNA Technologies, Inc. All PNA and biotinylated PNA were synthesized at PNA Bio Inc. Sequences are summarized in **Supplementary Table.1**.

### iSAT screening platform

#### S150 preparation

iSAT assay was adapted from previous method. Briefly, *E. coli* MRE600 was grown to log phase in LB at 37 °C and harvested at 6500 rpm 15 min at 4 °C. The pellet was resuspended in Buffer A. lysed open using French press. The lysate was cleared by centrifuge at 30,000 g for 30 min for twice. The supernatant was carefully layered over a 37.5% sucrose cushion and centrifuge at 90,000 g for 18 hours. The pellet (mainly contains 70S ribosomes) was resuspended in Buffer C (10 mM Tris-HOAc, pH 7.5 @ 4 °C, 60 mM NH_4_Cl, 7.5 mM Mg(OAc)_2_, 0.5 mM EDTA, 2 mM DTT) and saved for later r-protein purification, while the supernatant was centrifuged at 150,000 g for 4 hours. The upper 2/3 of the supernatant S150 extract was dialyzed against AED buffer AED three times over night and concentrated using Amicon (MWCO 3K) to A260 over 25. The concentrated S150 extract was then pelleted at 4,000 g to remove any precipitation. The resulted supernatant S150 extract was aliquoted, frozen in liquid nitrogen and store at -80 °C before usage.

#### r-protein purification

The frozen purified ribosome was thawed on ice and then diluted to 250-300 total A260 unit before adding spermine and spermidine to 0.2 mM and 2 mM respectively. 0.1 volume of 1 M Mg(OAc)_2_ was added to diluted ribosome before precipitation of rRNA with 2X volume of glacial acetic acid for 45 min. The resulted rRNA precipitation was removed by 30 min centrifugation at 16,000 g. The r-protein in the supernatant was then precipitated by 5 volumes of acetone at -20 °C overnight. The r-protein was pelleted at 10,000 g for 30 minutes. The resulted pelleted was dissolved in TP70 Urea buffer (HEPES pH 7.6 at room temperature, 10 mM Mg(Glu)_2_, 200 mM KGlu, 0.5 mM EDTA, 1 mM Spermidine, 1 mM Putrescine, 2 mM DTT, 6 M Urea) and dialyzed against 100 volume TP70 Urea buffer in Tube-O-Dialyzer™(G-Biosciences) at 4 °C overnight and then dialyzed against 100 volume TP70 buffer (TP70 Urea buffer without Urea) 90 min for three times at 4 °C. The TP70 concentration was measured by nanodrop for absorbance at 230 nm for concentration determination (4.17 µM/A230). The r-protein was aliquoted, frozen in liquid nitrogen and store at -80 °C before usage.

#### iSAT reaction

the iSAT reaction was performed in 57 mM Hepes-KOH, 1.5 mM spermidine, 1 mM putrescine, 10 mM Mg(Glu)_2_, and 150 mM KGlu at pH 7.5 with 2 mM DTT, 0.33 mM NAD, 0.27 mM CoA, 4 mM sodium oxalate, 2% w/v PEG-6000, 2 mM amino acids (Sigma-Aldrich), 1 nM pY71sfGFP plasmid encoding sfGFP, 0.1 mg/mL in-house made T7 RNA polymerase, 42 mM phosphoenolpyruvate (Roche, pH adjusted to 8.0 with KOH before addition) and NTP (all from sigma, pH to 8.0 with KOH, 1.6 mM ATP, 1.15 mM of GTP, CTP and UTP each), 45.3 μg/μL tRNA from E. coli MRE 600 (Roche), 227.5 μg/μL Folinic acid. The above components were premixed. A 5.36 µL aliquot of the premix was pipetted into 5 µl of S150 extract. Ribosomal proteins were added to a final concentration of 0.6 µM. iSAT reactions started after addition of plasmid encoding *rrnB* under T7 promoter at 4 nM and the reaction was immediately incubated on pre-warmed CFX Connect Real-time System (Bio-Rad) at 37 °C for 90 mins in 96 well plates (Applied Biosystems). sfGFP production was detected by fluorescence measurement at 2 min intervals (excitation: 450-490 nm, emission: 510-530 nm). For ASO added samples, the ASO was heated at 95 °C for 5 min and added into the reaction before addition of *rrnB* plasmid. The lag time is obtained from the Bio-Rad base line fitting and subtraction and compared with control iSAT.

#### RNA target rescue

In target rescue experiment, 1.5-fold RNA fragment was pre-annealed with PNA/LNA at 95 °C for 5 min before it was added into a normal iSAT reaction with final PNA/LNA concentration at 10 µM.

#### Z score statistics

For DNA library screening, the Z-score is calculated for Δ*lag* and Δ*F*. For PNA screening at various concentration, the Z-score is only calculated for Δ*F*. In both scenarios the following formula was used:

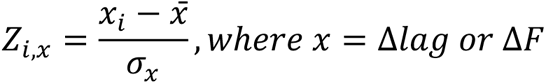

### Sucrose gradient fractionation

For sample preparation, 10 iSAT reactions were pooled and mixed with 250 uL 3X non-dissociation gradient buffer. The mixture was loaded onto a 11 mL 10-40 % w/v sucrose gradient (50 mM Tris-HCl 7.8, 100 mM NH_4_Cl, 10 mM MgCl_2_, 6 mM β-mercaptoethanol) then centrifuged in a Beckman SW41 rotor at 26,000 rpm for 16 hours at 4 °C. Gradients were fractionated using a Brandel gradient fractionator with flow rate set at 1.5 mL/min. Based on the UV 254 nm trace, gradient fractions between 30S and 70S were collected and combined. To prepare the fractions for cryo-EM analysis, 3X volumes of buffer A were added prior to concentration in a 100 kDa cutoff concentrator (Amicon) 3 times to eliminate sucrose and to equilibrate to buffer A.

### Cy5-Streptavidin blot

After gradient separation, the fractions were diluted in TBST and filtered through Hydrobond-N^+^ membrane with Dot-Blot System (Schleicher & Schulell) under vacuum. The membrane was blocked with 0.5% bovine serine albumin in TBST for 30 min, then blotted against 1:1000 Cy5-Streptavin (Life Technologies) at RT for 1 hour with gentle shaking. The membrane was washed 4 times with TBST before imaged with VersaDoc using Cy5 channel.

### Grid preparation for data collection

Grids were prepared for a total of 7 samples-control, RA20, RB3248, RB2937, RB2860, RB2737 and RB0811. All samples were in non-dissociation gradient buffer. For all samples, Au-Flat R1.2/1.3 grids (Protochips) were plasma treated using Solarus Plasma Cleaner 950 (Gatan, 75% Argon, 25% Oxygen) for 10 (control and RB3248) or 15 seconds (rest of the samples). For all samples, 3 µL of ∼0.1 µM was applied to plasma treated grids, manually blotted with a Whatman filter paper grade 1 (Cytiva) to wick excess sample and immediately plunge frozen in liquid ethane using a custom homemade device. Vitrification for all samples was performed in a cold room maintained at 4°C with a humidity of greater than 80%.

### Data collection

Data was collected on a FEI Titan Krios (Thermo Fisher) equipped with either a K2 Summit (control and RB3248) or K3 BioQuantum with CDS mode (rest of the samples) direct electron detector with energy filter slit width of 20 eV (all samples). For all samples, automated data collection was performed using the Leginon software package^32^ with a pixel size of 1.05 Å per pixel (control and RB3248) or 0.83 Å per pixel (rest of the samples). Movies were recorded with an exposure time of 4.4 seconds (control and RB3248) or 2 seconds (rest of the samples), with 55 frames per movie (control and RB3248) or 80 frames per movie (rest of the samples) and with a dose rate of 6.32 e-/pixel/second and total dose of 25.2 e-/Å (control and RB3248) or 12.645 e-/pixel/second and total dose of 36.71 e-/Å (RB2937 and RB0811) or 8.75 e-/pixel/second and total dose of 25.4 e-/Å (rest of the samples). Data was collected with a 20° stage tilt (control and RB3248) or 30° stage tilt (RB2937 and RB0811) or no stage tilt (rest of the samples).

### Heterogeneous reconstruction of PNA-intermediates

The raw movies were directly imported into CryoSPARC^33^ and processed by MoionCor2^34^ followed by CTFFIND4^35^. Control sample was picked with a template generated by a low-resolution C-class with diameter set to 300 Å. After two rounds of 2D classification two remove junk particles. The intermediate-like particles were subjected to ab-initio reconstruction, requesting for 4 classes using default parameters. Each resulting interpretable class was subjected to another round of ab-initio reconstruction using the same parameters. This procedure was performed iteratively until the particle number in a class was less than 2,000, in which case the 3D reconstruction would result in low resolution maps as described in previous method. All reconstructions with fewer than 2,000 particles were subjected to ab-initio reconstruction, requesting 1 class prior to 3D refinement in CryoSPARC. All refined density maps were aligned and resampled to a same 50S ribosome reference (bL17-depletion dataset E) in ChimeraX^36^. All maps with resolution below 10 Å were discarded and the remaining resampled density maps were thresholded at intensity 1.00. Pairwise difference maps were calculated for the binarized (thresholded at 0.025) maps the sum of the difference map A-B and difference map B-A for hierarchical clustering using the Ward linkage. Maps were displayed with the resulting dendrogram, and pairs of maps with a difference of <10 KDa were merged into one class. The value of 10 kDa was defined previously as a valid merging criterion, as it represents the average molecular weight of all proteins and rRNA helices constituting the LSU. This step is important as similar classes can emerge from hiding at various stages of the iterative subclassification. The merged particles were next subjected to an ab-initio reconstruction and 3D refinement to produce the final map for the class.

In other datasets, particles were picked by blob picker with Diameter set to 200 to 300 Å and extracted with fourier crop to same box size to the control with 1.05 Å as pixel size. After several rounds of 2D classification to clean out the junk particles. The intermediate-like particles were mixed with control particles for first round of ab-initio classification. The control particles in intermediate-like classes were then removed using particle set tool. This process helped with alignment of rare intermediate classes that might be otherwise hard to align in the first ab-initio reconstruction. After separation, the iterative subclassification process was the same as control sample. The particle and volume metadata were summarized in **Supplementary Table.2**.

Finally, was aligned to reference map, a 10 Å density map of 50S in 4YBB, and scaled according to 4YBB map and binarized at 3.5 sigma for each map using in-house script. The final hierarchical clustering analysis was then performed on resulted binarized scaled maps across seven datasets with previous maps solved in bL17-depletion, Δ*deaD* and Δ*srmB* dataset in the same way to allow comparison.

### 3D Flex reconstruction and modeling of RA20-B-c Class

From one iterative subclassification, RA20-B-c1 and RA20-B-c2 were split. The particles before the subdivision were used for an ab-initio reconstruction with one class. The class was refined by homogeneous refinement to 4.71 Å with fuzzy density at the mis-docked domain III/VI region. The consensus reconstruction was then subjected to 3D Flex training. The resolution reported in 3D Flex Reconstruction was 5.27 Å, but with more solid density in the same region. The model was built into the density with Coot using 4YBB as initial model. The movies were generated with Flex generate in CryoSPARC.

### Contact dependency map

The Occupancy of each assembly block was calculated using previous pipeline with slight modification. For structure elements, 50S in 4YBB model was used after align to the volume sum of the whole dataset. Different individual PDB and volume files are generated through in-house script. Briefly, volume for each defined structure block in **Supplementary Material 1** was generated using molmap in ChimeraX with 7 Å resolution and binarized with 0.01 threshold. For each of the density maps thresholded at intensity 0.035. The number of voxels above threshold are counted in each structure element then normalized to the total number of voxels in the structure element. The occupied fraction for each block in each density map is then normalized to uL24 if it is a protein or otherwise to H2.

The dependency between any pair of blocks (*i,j*) was obtained by quadrant analysis of a scatter plot of the occupancy for block *i* on the x-axis and block *j* on the y-axis. The dashed binarization lines for the horizontal and vertical directions were calculated by the following equation (eq.1),

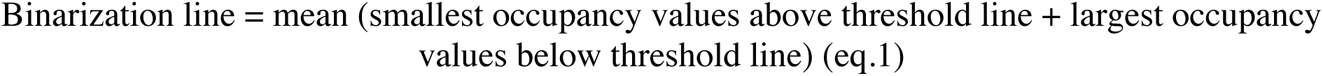

* threshold is defined in **Supplementary Material 2.**

The x and y binarization lines divide the scatter plot into 4 quadrants: Q1 = lower left, Q2 = lower right, Q3 = upper left, Q4 = upper right. To infer the relationship between block *i* and *j*, the number of points in each quadrant were counted for the scatter plot. The relationship between block *i* and *j* falls into one of three scenarios. With points only in Q1 and/or Q4, blocks *i* and *j* are correlated. With dots in both Q2/Q3, blocks *i* and *j* are not correlated. With dots only in Q1/Q2/Q4 or Q2/Q4, block *j* should depend on block *j*. With dots in only Q1/Q3/Q4 or Q3/Q4, block *i* should depend on block *j*. If block *i* depends on block *j*, an arrow from *j* to *i* will be drawn in the dependency map.

The comprehensive dependency plot is now ready for pruning with networkx package ^37^. First, the dependency between two structure elements without any contacts will be removed. The non-direct dependency arrows were then further pruned. For example, arrow from 1 to 3 will be removed, if 1 to 2 and 2 to 3 both exist. the pairwise contacts for each structure elements are calculated with “Structure Analysis – Contact” in ChimeraX using in-house script.

### Code Accessibility

The code for volume curation (alignment, resample and rescale), hierarchical analysis, contact analysis, occupancy analysis and quadrant-dependency analysis can be provided upon request.

## Acknowledgements.

This work was supported by a grant from the NIH GM-136412 (to J.R.W) and NIH U54 AI170855 and the Hearst Foundations Developmental Chair (to D.L.).

## Data availability

The density maps have been deposited in the EMDB with the following accession numbers: ctrl-C-a1: EMD-44483, ctrl-C-a2: EMD-44485, ctrl-C-b1: EMD-44487, ctrl-C-b2: EMD-44488, ctrl-C-b3: EMD-44564, ctrl-E-a1: EMD-44565, ctrl-E-a3: EMD-44566, ctrl-E-a2: EMD-44567, ctrl-E-a4: EMD-44568, RA20-B-a1: EMD-44569, RA20-B-a2: EMD-44570, RA20-B-c1: EMD-44571, RA20-B-c2: EMD-44572, RA20-C-a1: EMD-44573, RA20-C-b1: EMD-44574, RA20-E-a1: EMD-44575, RB0811-C-a1: EMD-44576, RB0811-C-b1: EMD-44577, RB0811-C-b2: EMD-44578, RB0811-E-a1: EMD-44579, RB0811-E-a2: EMD-44580, RB0811-E-a3: EMD-44581, RB0811-E-a4: EMD-44582, RB2737-C-a1: EMD-44583, RB2737-C-a2: EMD-44604, RB2737-C-a3: EMD-44605, RB2737-E-a1: EMD-44606, RB2737-E-a2: EMD-44607, RB2737-G1: EMD-44608, RB2737-G2: EMD-44609, RB2860-C-a1: EMD-44610, RB2860-E-a1: EMD-44610, RB2937-C-a1: EMD-44612, RB2937-C-a2: EMD-44613, RB2937-C-b1: EMD-44614, RB2937-E-a1: EMD-44615, RB2937-E-a2: EMD-44616, RB2937-E-a3: EMD-44617, RB2937-E-a4: EMD-44618, RB3248-C-a1: EMD-44619, RB3248-C-b1: EMD-44620, RB3248-E-a1: EMD-44621, RB3248-E-a2: EMD-44622, RB3248-E-b1: EMD-44623, RB3248-J1: EMD-44624, RB3248-J2: EMD-44625, RB3248-J3: EMD-44626.

