## Extended Data Figure and Supplementary Table for "Domain consolidation in Bacterial 50S assembly revealed by Anti-Sense Oligonucleotide Probing"

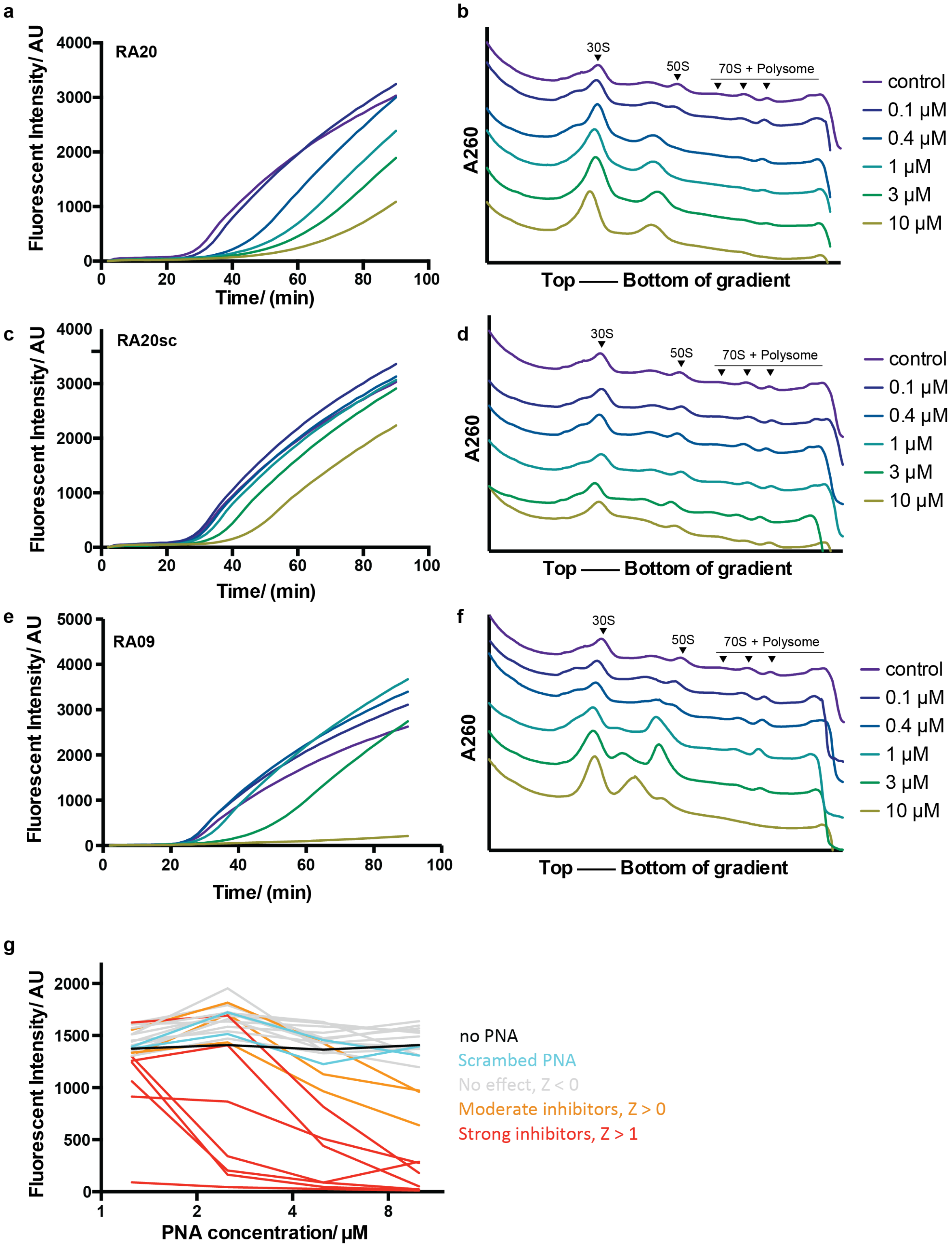


**Extended Data Fig.1 |** Titration of PNA in iSAT assay

**Extended Data Fig.1 (continued)** Fluorescent traces for iSAT reaction with RA20 (**a**), scrambled RA20 (RA20sc) (**c**) and RA09 (**e**) and corresponding sucrose gradient profiles (**b d f**). The PNAs were titrated from 10 μM to 0.1 μM with three-fold dilution. The 30S, 50S, 70S and polysome peaks were marked on the gradient profile for the control sample. The traces in **a-f** are colored by the concentration of the PNA according to the rightmost color legends. **g**) End-point fluorescence for PNA library at 10, 5, 2.5 and 1.25 μM. The curves are colored according to the *Z*-scores in **Fig.1 h**.

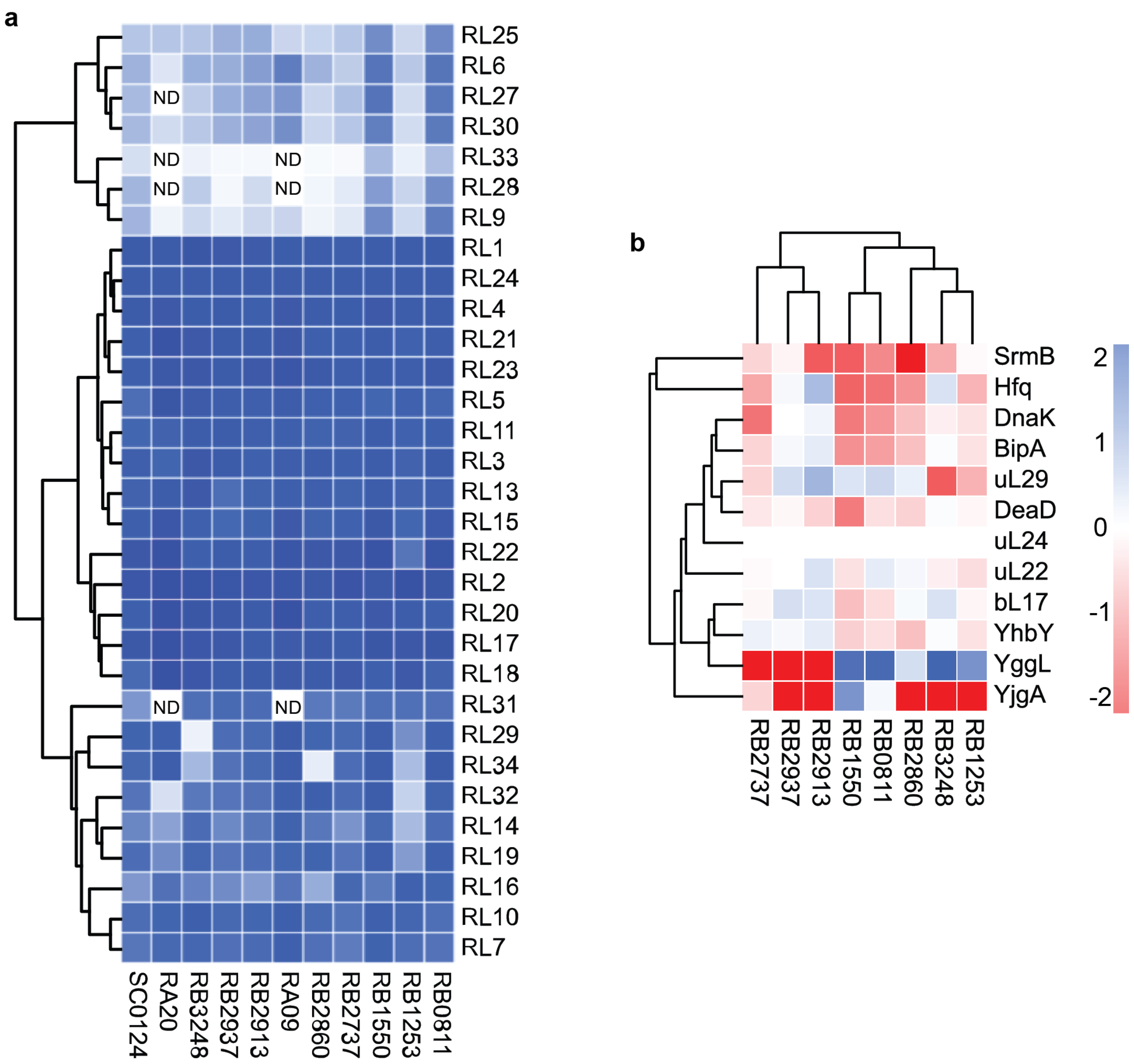


**Extended Data Fig.2 |** Proteomics analysis of intermediate fractions

**Extended Data Fig.2 (continued) a**) Heatmap representation for quantitative proteomics analysis for r-proteins. The abundance was normalized to N15 samples then uL24 abundance in each sample. (Blue:1, white: 0). ND means there is no peptide detected in either N14 or N15 peptide search. **b**) Peptide spectrum matches (PSMs) for selected assembly factors. The PSMs were normalized to total N14 PSMs then control sample. The color scheme from red to blue represents the $\log_2$ fold change ($\log_2$FC).

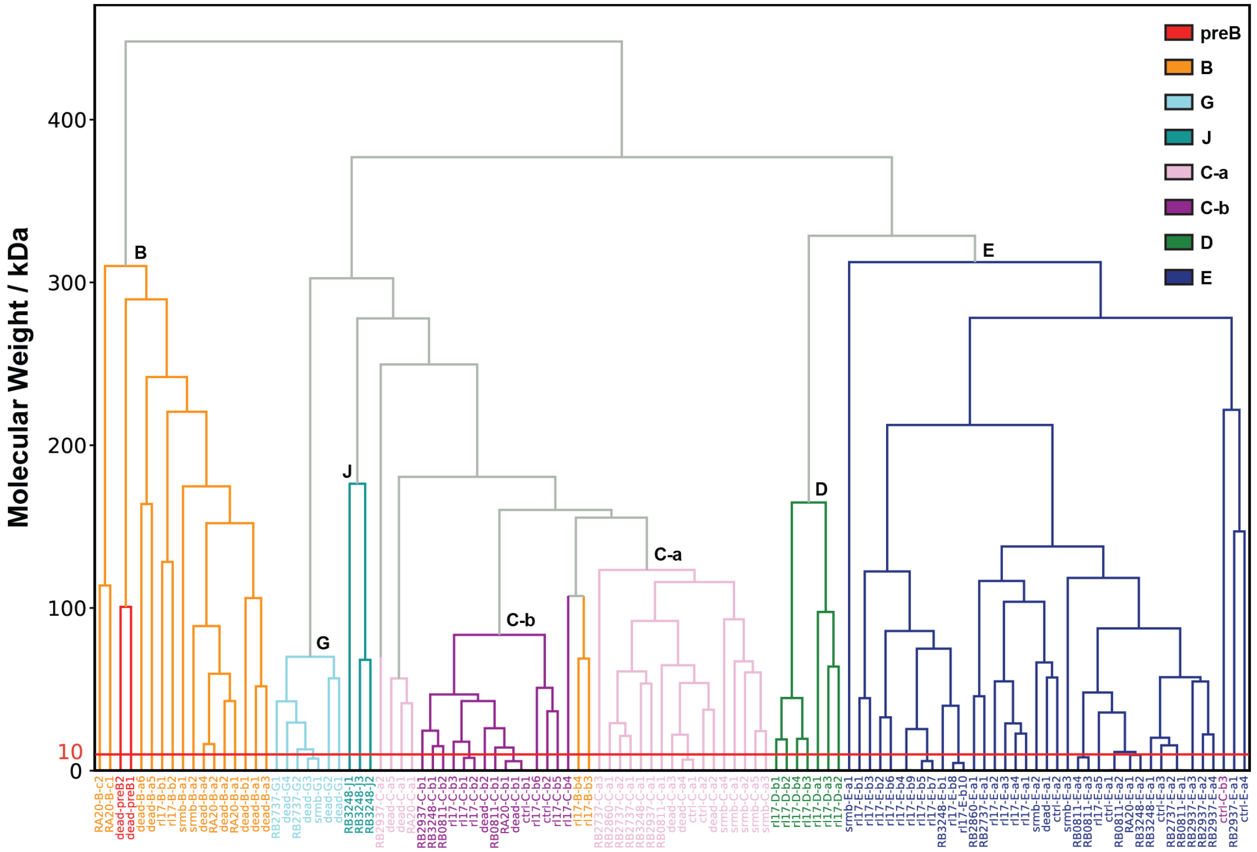


**Extended Data Fig.3 |** Comparison with previous classes

The Euclidean distance matrix, based on molecular weight in kDa, was calculated among 47 density maps and previous density maps from bL17-depletion, Δ*deaD* and Δ*srmB* datasets. The dendrogram resulting from hierarchical clustering is displayed, with the 8 main class branches colored accordingly.

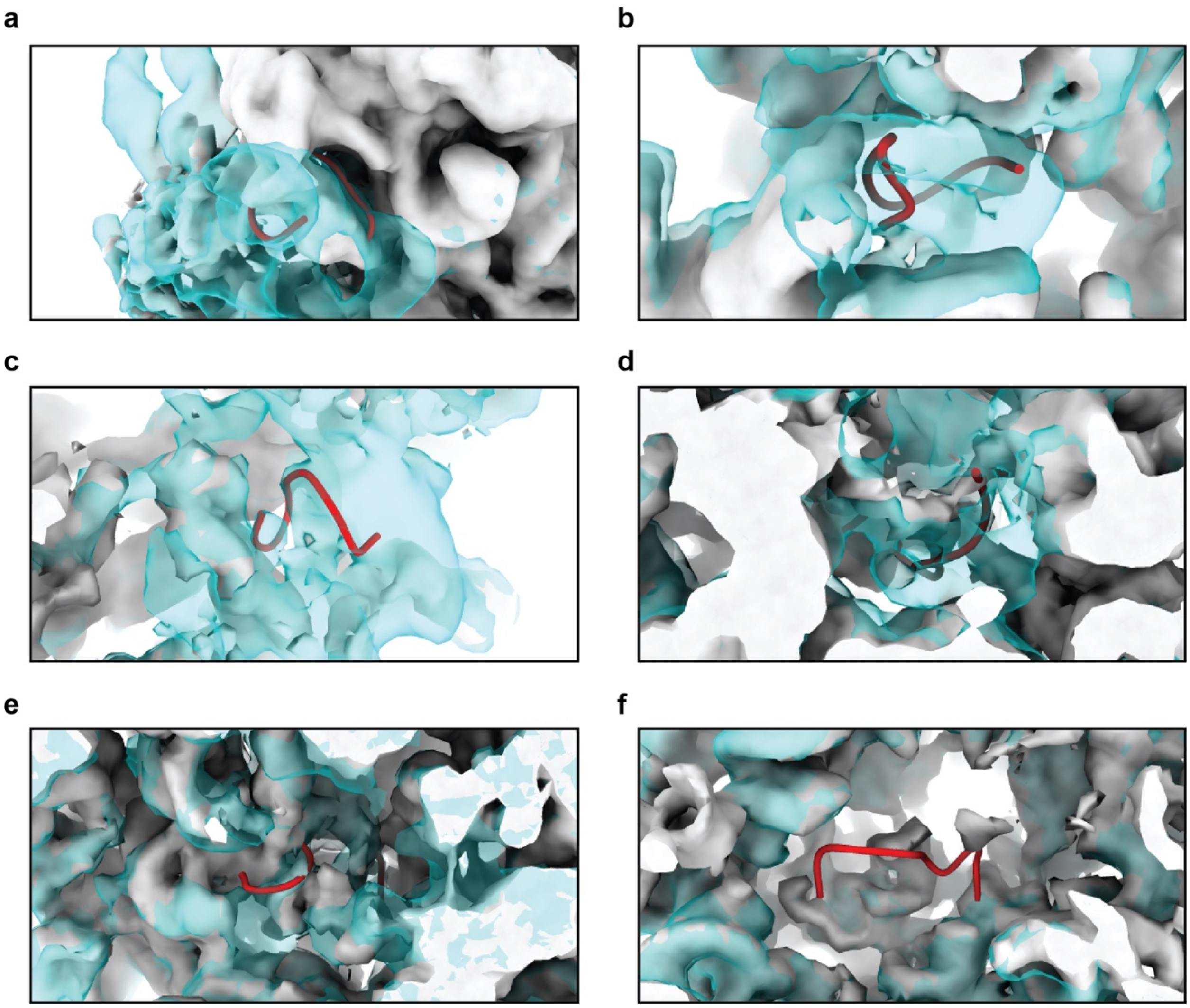


**Extended Data Fig.4 |** PNA targeting sites in control average map

**a-g**) shows the PNA targeting site (red ribbons) on corresponding average map (from **a** to **g**: RA20, RB3248, RB2937, RB2860, RB2737 and RB0811, shown in light grey density). The control average map was shown in transparent cyan density.

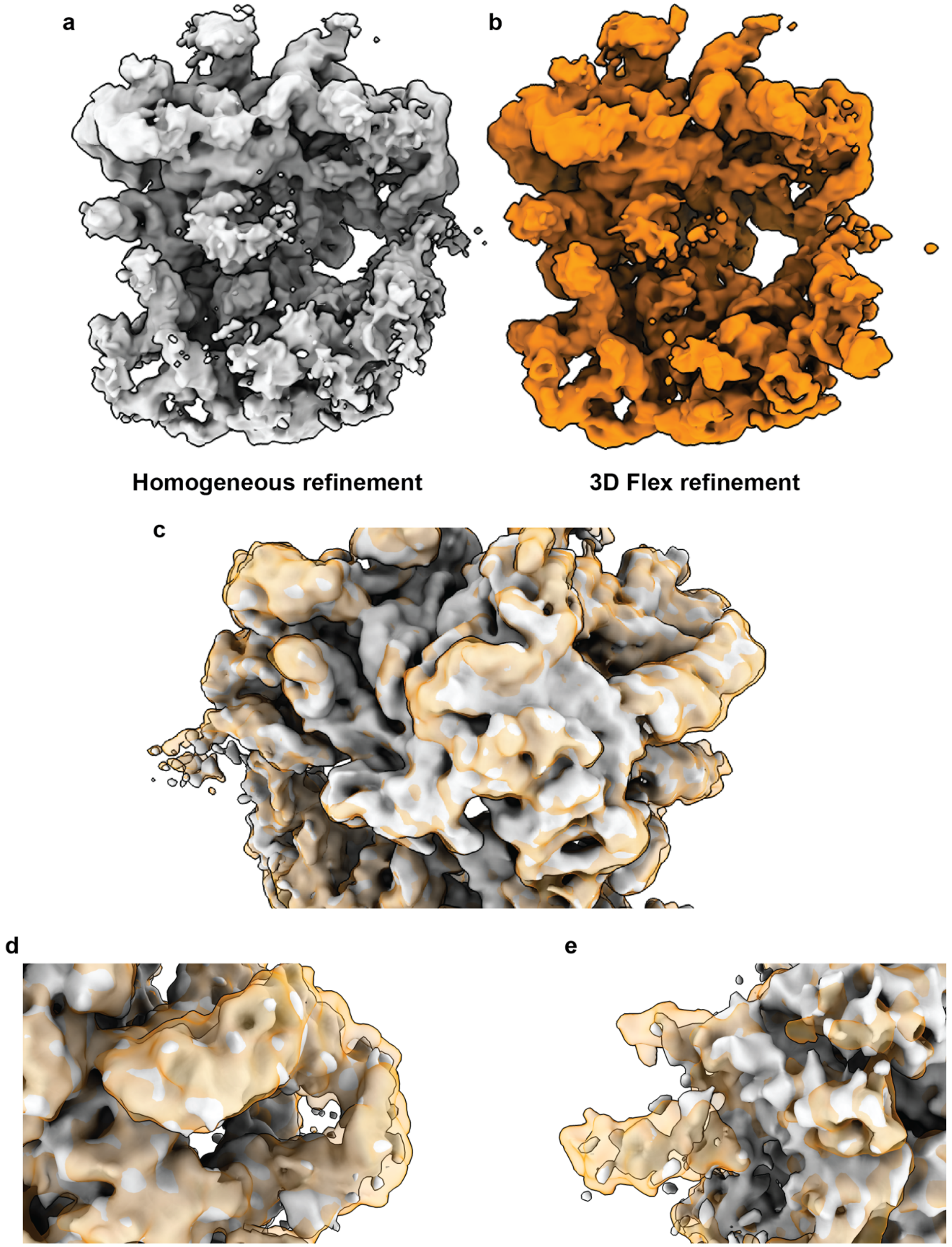


**Extended Data Fig.5 |** Comparison between density map from homogeneous refinement and from 3D Flex refinement for RA20-B-c class

**Extended Data Fig.5 (continued)** The homogeneous refinement and 3D flex refinement map for RA20-B-c class are shown in **a**) and **b**) respectively. Zoomed in comparison for **c**) domain I/II, **d**) domain III, and **e**) domain VI are shown with map from 3D flex refinement in orange transparent volume and map from homogeneous refinement in light grey.

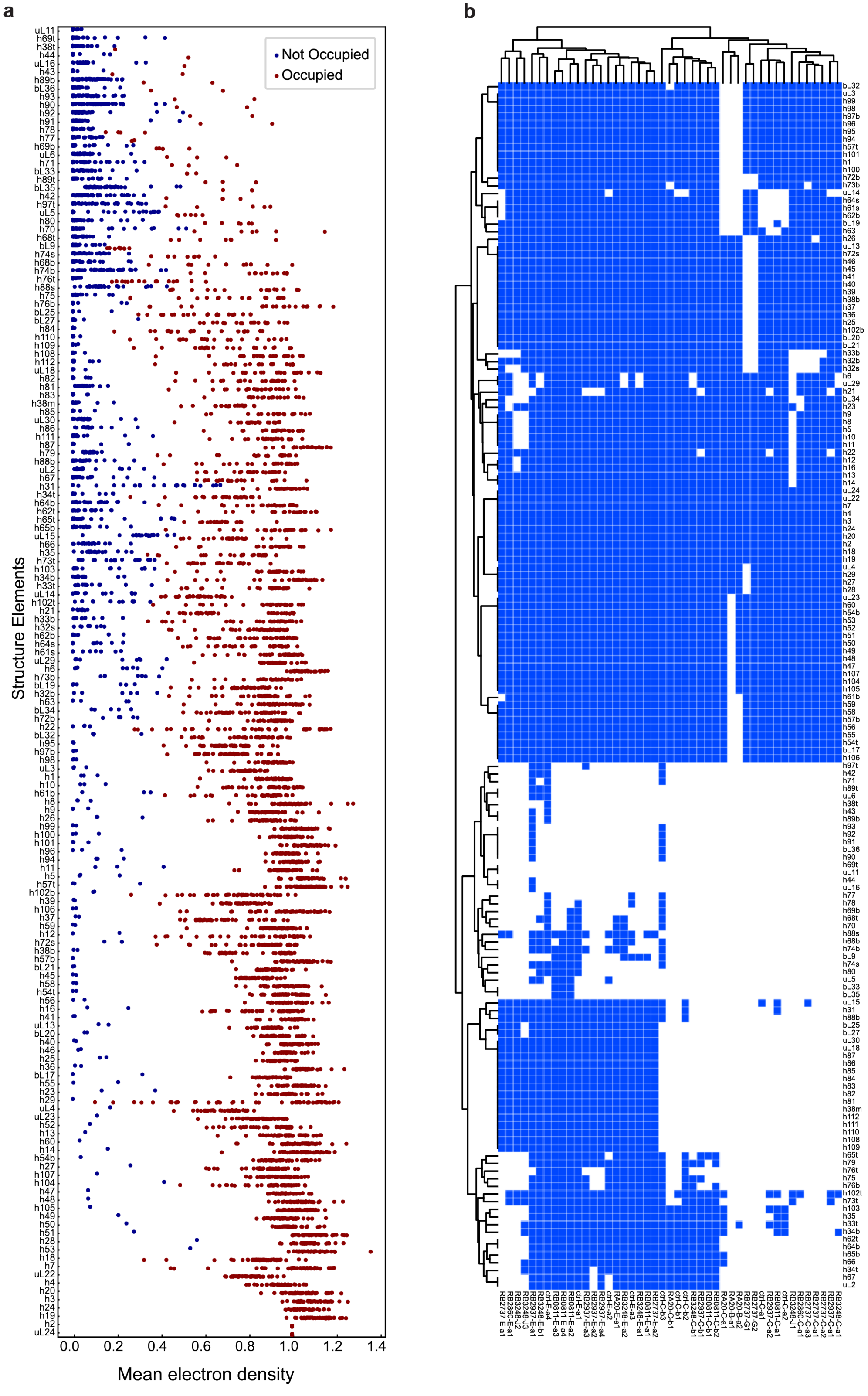


**Extended Data Fig.6 |** Generation of Binarized Occupancy Matrix

**Extended Data Fig.6 (continued) a**) Strip plot for each structure elements. There are 157 structure elements on the y-axis. Each dot in one strip represents the occupancy (x-axis) for one structure element in one electron density map. The non-occupied dots were colored blue, while the occupied dots were colored red. b) Occupancy of 45 intermediate density maps (without RA20-B-c classes) from PNA-inhibited dataset in terms of the 157 structure elements, binarized with threshold in **a**) and used for dependency analysis.

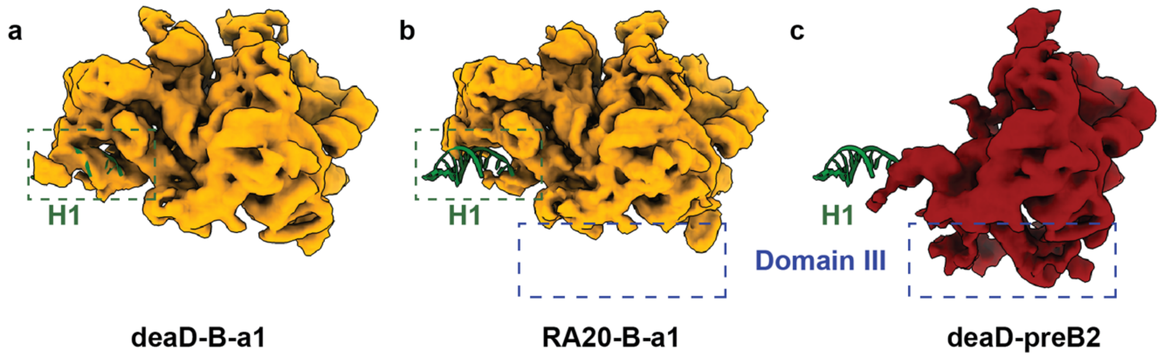


**Extended Data Fig.7 |** Missing of H1 density in different classes

Back view of **a**) deaD-B-a1, **b**) RA20-B-a1 and **c**) deaD-preB2 with atomic model of H1 helix (green), highlighted in green boxes. The empty and partial domain III density in b) and c) were highlighted in blue boxes.

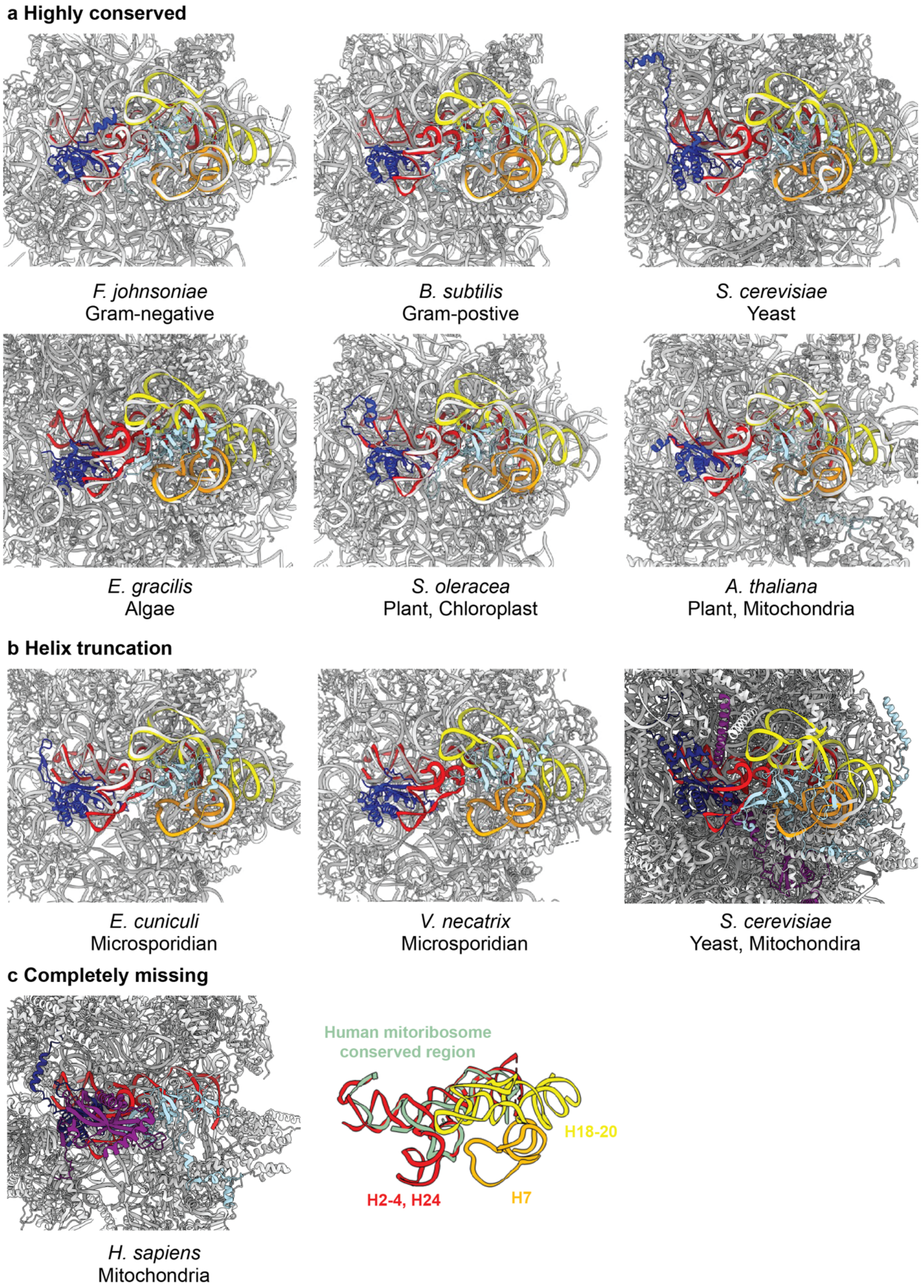


**Extended Data Fig.8 |** Smallest consensus core in different species

**Extended Data Fig.8 (continued)** The LSU models for different species are shown and classified according to the similarities in helix structure, **a**) highly conserved, **b**) truncated and **c**) completely missing. The rRNA helices in *E. coli* smallest consensus core identified in this work is shown in colorful ribbons (red: H2-4, H24, orange: H7, yellow: H18-20). The LSU rRNA in other species are shown in light grey. The uL22 and uL24 homologs for *E. coli* in other species are shown in dark blue and light blue respectively. The uL23 in yeast mitoribosome and mL45 in human mitoribosome were colored purple for easy visualization. The PDB IDs are (in sequential order): 7JIL, 6HA1, 5JCS, 6ZJ3, 5X8P, 6XYW, 7QEP, 6RM3, 5MRC, 7QI4. For human mitoribosome, the rRNA homologous regions are colored green in the right panel and overlaid with H2-4, H24, H7 and H18-20 in *E. coli*. The secondary structures were drawn using RiboVision2.

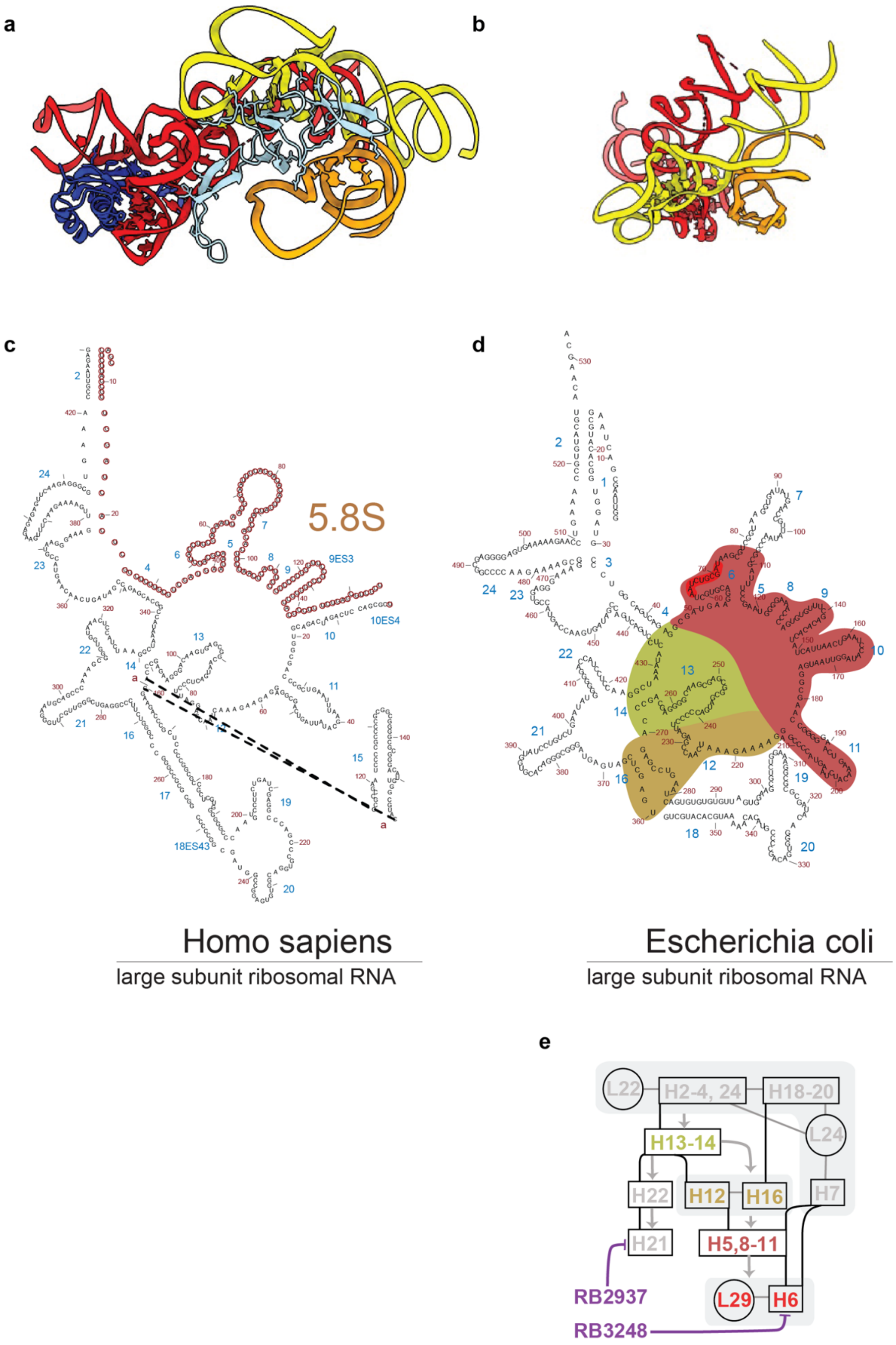


**Extended Data Fig. 9 |** Smallest consensus core and 5.8S

**Extended Data Fig. 9 (continued)**

**a**) Back and **b**) side views of the smallest consensus core. Secondary structure for part of domain I rRNA in **c**) *H. sapiens* and **d**) *E. coli*. The 5.8S in *H. sapiens* is colored brown. The helix defect in *E. coli* LSU with RB3248 are marked with according to dependency in **e**).

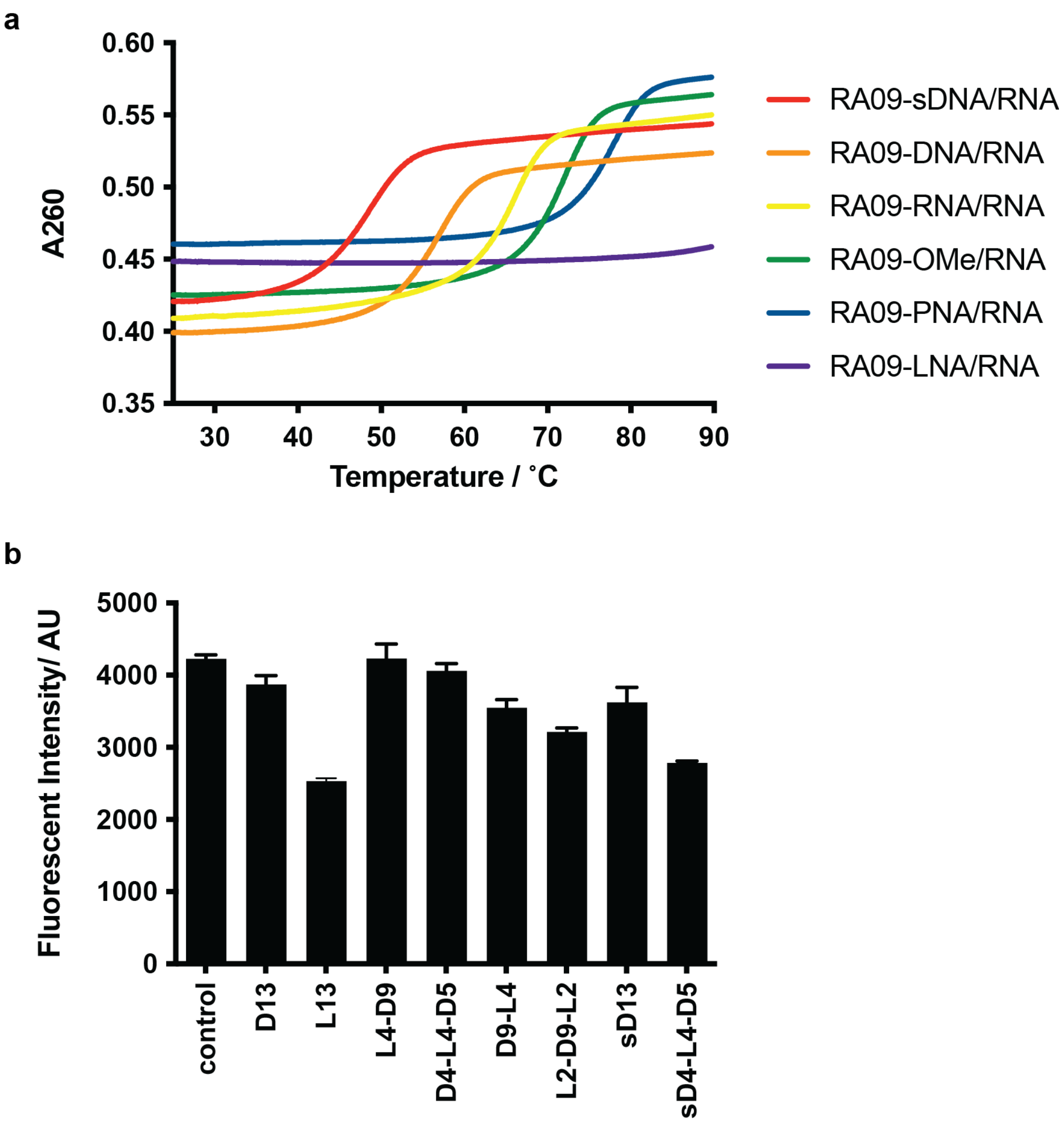


**Extended Data Fig. 10 |** Properties of ASO analogs

**a**) Melting curves of ASO analogs of RA09 to the RNA target were measured at 3 µM in iSAT buffer. The temperature ranges from 25 to 85 ˚C and during the temperature ramping, the A260 nm was measured. **b**) End-point fluorescence of RA20 with different locked site and phosphorothioate (PS) backbones. L*m*D*n* means the locked state starting from 5' end of the ASO. For example, L4D9, means first four ribose for RNA20 are 2'-5' locked, while the rest of it remained DNA backbone. "s" denotes the backbone of the ASO is replaced with PS.

**Supplementary table**

**Supplementary Table.1a Sequences for 1st library**

| *Name* | Sequence | Length | GC |
|---|---|---|---|
| **RA01** | GTTTGACGCTCAAAGAATT | 19 | 0.37 |
| **RA03** | CCTCTACGAGACTCAA | 16 | 0.50 |
| **RA04** | GGAGGTGATCCAACC | 15 | 0.60 |
| **RA05** | CACTACAAAGTACGCTTCT | 19 | 0.42 |
| **RA06** | ATTGCTTATCACGCGT | 16 | 0.44 |
| **RA07** | CAAACCAGCAAGTGGCGTCC | 20 | 0.60 |
| **RA08** | CTTAACCTCACAACCCGA | 18 | 0.50 |
| **RA09** | GGAGTATTTAGCCTTG | 16 | 0.44 |
| **RA10** | CGAAACAGTGCTCTAC | 16 | 0.50 |
| **RA11** | GAACAGCCATACCCTT | 16 | 0.50 |
| **RA12** | GTAAGGTTAAGCCTCA | 16 | 0.44 |
| **RA13** | GCTACTGCCGCCAGGC | 16 | 0.75 |
| **RA14** | TCTAGACGAAGGGGAC | 16 | 0.56 |
| **RA15** | TTATCGTTACTTATG | 15 | 0.27 |
| **RA16** | GCGAGTTCAATTTC | 14 | 0.43 |
| **RA18** | TTCCATTCAGAACC | 14 | 0.43 |
| **RA19** | CGACATCGAG | 10 | 0.60 |
| **RA20** | TAGTCGCTTAACC | 13 | 0.46 |
| **RA21** | GTTACTCTTTA | 11 | 0.27 |
| **RA22** | ACAACCGTCG | 10 | 0.60 |
| **RA23** | CAGGAACCCTT | 11 | 0.55 |
| **RA24** | TTTCCCACTTAA | 12 | 0.33 |
| **RA25** | GGTTTGGGGTA | 11 | 0.55 |
| **RA26** | TTATAGTTACGGC | 13 | 0.38 |

**Supplementary Table.1b Sequences for 2nd library**

| *Name* | Sequence | Length | GC |
|---|---|---|---|
| **SC0124** | CGACATAGTCCAC | 13 | 0.54 |
| **SC0180** | GAGATCGATGCCC | 13 | 0.62 |
| **RB2263** | ACATCCTGGCTGT | 13 | 0.54 |
| **RB2206** | GCCGACTCGACCA | 13 | 0.69 |
| **RB2172** | TGCATGGTTTAGC | 13 | 0.46 |
| **RB1550** | TTGCAGCCAGCTG | 13 | 0.62 |
| **RB3248** | CTTATCGCAGATT | 13 | 0.38 |
| **RB3202** | CATTCGGAAATCG | 13 | 0.46 |
| **RB3152** | AACCTATGGATTC | 13 | 0.38 |
| **RB3000** | GTATCGCGCGCCT | 13 | 0.69 |
| **RB2937** | CGTGTCCCGCCCT | 13 | 0.77 |
| **RB2913** | CCCCCATATTCAG | 13 | 0.54 |
| **RB2860** | CGGTACTGGTTCA | 13 | 0.54 |
| **RB2789** | ACTGCTTGTACGT | 13 | 0.46 |
| **RB2744** | CCCATTATACAAA | 13 | 0.31 |
| **RB2737** | TCGCTGACCCATT | 13 | 0.54 |
| **RB1461** | CCCATCAATTAAC | 13 | 0.38 |
| **RB1401** | CCGTTATAGTTAC | 13 | 0.38 |
| **RB1253** | TCACGGGGTCTTT | 13 | 0.54 |
| **RB1200** | CACCTATCCTACA | 13 | 0.46 |
| **RB0811** | GAGCCGACATCGA | 13 | 0.62 |
| **RA09-PNA** | GGAGTATTTAGCCTTG | 16 | 0.44 |
| **RA20-PNA** | TAGTCGCTTAACC | 13 | 0.46 |
| **RA20sc-PNA** | ATTGGCTCATCAC | 13 | 0.46 |

**Supplementary Table.2 Metadata for electron density maps**

| *Name* | *particle number* | *resolution/ Å* | *EMD* |
|---|---|---|---|
| ***ctrl-C-a1*** | 2120 | 5.8 | EMD-44483 |
| ***ctrl-C-a2*** | 1910 | 6.2 | EMD-44485 |
| ***ctrl-C-b1*** | 5675 | 4.7 | EMD-44487 |
| ***ctrl-C-b2*** | 3672 | 4.8 | EMD-44488 |
| ***ctrl-C-b3*** | 1633 | 6.0 | EMD-44564 |
| ***ctrl-E-a1*** | 5153 | 4.3 | EMD-44565 |
| ***ctrl-E-a3*** | 1732 | 5.1 | EMD-44566 |
| ***ctrl-E-a2*** | 1631 | 6.1 | EMD-44567 |
| ***ctrl-E-a4*** | 1006 | 6.3 | EMD-44568 |
| ***RA20-B-a1*** | 3022 | 4.2 | EMD-44569 |
| ***RA20-B-a2*** | 4218 | 3.9 | EMD-44570 |
| ***RA20-B-c1*** | 3904 | 5.2 | EMD-44571 |
| ***RA20-B-c2*** | 3608 | 6.1 | EMD-44572 |
| ***RA20-C-a1*** | 2201 | 4.4 | EMD-44573 |
| ***RA20-C-b1*** | 2754 | 4.6 | EMD-44574 |
| ***RA20-E-a1*** | 1898 | 4.2 | EMD-44575 |
| ***RB0811-C-a1*** | 1707 | 6.1 | EMD-44576 |
| ***RB0811-C-b1*** | 1457 | 5.9 | EMD-44578 |
| ***RB0811-C-b2*** | 1430 | 6.1 | EMD-44579 |
| ***RB0811-E-a1*** | 7066 | 4.6 | EMD-44580 |
| ***RB0811-E-a2*** | 3430 | 4.7 | EMD-44584 |
| ***RB0811-E-a3*** | 1857 | 5.1 | EMD-44581 |
| ***RB0811-E-a4*** | 1236 | 5.5 | EMD-44582 |
| ***RB2737-C-a1*** | 2743 | 4.4 | EMD-44583 |
| ***RB2737-C-a2*** | 2558 | 4.7 | EMD-44604 |
| ***RB2737-C-a3*** | 1979 | 5.3 | EMD-44605 |

| | | | |
|---|---|---|---|
| ***RB2737-E-a1*** | 6682 | 4.2 | EMD-44606 |
| ***RB2737-E-a2*** | 2195 | 4.8 | EMD-44607 |
| ***RB2737-G1*** | 2437 | 4.7 | EMD-44608 |
| ***RB2737-G2*** | 2468 | 4.9 | EMD-44609 |
| ***RB2860-C-a1*** | 13429 | 4.3 | EMD-44610 |
| ***RB2860-E-a1*** | 3174 | 5.5 | EMD-44611 |
| ***RB2937-C-a1*** | 1661 | 6.9 | EMD-44612 |
| ***RB2937-C-a2*** | 1418 | 7.5 | EMD-44613 |
| ***RB2937-C-b1*** | 5244 | 5.1 | EMD-44614 |
| ***RB2937-E-a1*** | 2568 | 5.9 | EMD-44615 |
| ***RB2937-E-a2*** | 2507 | 6.2 | EMD-44616 |
| ***RB2937-E-a3*** | 1451 | 6.3 | EMD-44617 |
| ***RB2937-E-a4*** | 7355 | 5.0 | EMD-44618 |
| ***RB3248-C-a1*** | 1811 | 6.5 | EMD-44619 |
| ***RB3248-C-b1*** | 1942 | 5.1 | EMD-44620 |
| ***RB3248-E-a1*** | 2362 | 4.9 | EMD-44621 |
| ***RB3248-E-a2*** | 1983 | 5.0 | EMD-44622 |
| ***RB3248-E-b1*** | 1796 | 4.9 | EMD-44623 |
| ***RB3248-J1*** | 5827 | 4.2 | EMD-44624 |
| ***RB3248-J2*** | 2415 | 5.6 | EMD-44625 |
| ***RB3248-J3*** | 2461 | 5.3 | EMD-44626 |
